# Brain Structure-Function Association as a Compensatory Factor in Cognitive Aging

**DOI:** 10.64898/2026.07.16.739039

**Authors:** Tzu-Chen Lung, David A. Hoagey, Karen M. Rodrigue, Michael D. Rugg, Kristen M. Kennedy

## Abstract

Functional activity in response to increasing task difficulty (BOLD-modulation) in regions of the default mode network (DMN/task-negative) and multiple demand network (MDN/task-positive) shows age-related decline, which is associated with reduced cognitive performance. While BOLD-modulation in the DMN begins to decline in middle-age, white matter structural connectivity declines earlier in MDN regions. We conjecture that to maintain executive function (EF) performance with aging, altered BOLD-modulation in DMN during task engagement serves as a compensatory reaction to structural decline in MDN. To test this possibility, functionally-guided tractography was applied in 160 healthy adults aged 20-94 to locate white matter connecting MDN and/or DMN regions that were active during a distance judgement paradigm. Specifically, analyses examined the effects of 1) age on white matter tracts (fractional anisotropy; FA); 2) structure-function association between FA and BOLD-modulation; and 3) age, positive/negative BOLD-modulation, and quadratic FA on EF. Age-related decline was found in one MDN tract (U-shaped tract) and a significant structure-function association was found in inferior fronto-occipital fasciculus (IFOF). Additionally, an interaction effect between quadratic frontal-insular (U-shaped) tract FA, age, and positive/negative BOLD-modulation was found for inhibition performance, supporting the hypothesis of compensatory functional activity effects on executive function for older adults with altered white matter microstructure.

## 1. INTRODUCTION

Age-related decline of structural integrity and functional activity in the brain contributes to declines in working memory and executive functions (Brown et al., 2019; Hakun et al., 2015; Webb et al., 2019, 2020; Zhu et al., 2015). Functional neuroimaging studies have established that two opposing largescale, intrinsic brain networks contribute to higher-order cognitive functions such as executive functions and working memory. The multiple demand network (MDN) comprises brain regions that demonstrate task-related positive blood oxygenation-level dependent (BOLD) activity, which increases as task difficulty increases. In contrast, the default mode network (DMN) demonstrates task-related negative BOLD activity, which becomes more negative-going as task difficulty increases (Grady et al., 2010). Across the adult lifespan, BOLD modulation to task difficulty weakens with older age in both networks and has been associated with decreased working memory (Kennedy et al., 2017; Rieck et al., 2017). The joint contribution (or coupling) of positive BOLD modulation (PosMod) and negative BOLD modulation (NegMod) has been linked to better cognitive control (Spreng et al., 2010; Spreng et al., 2016; Turner & Spreng, 2015), and higher fluid intelligence (Rieck et al., 2017).

Although age-related decreases in PosMod and NegMod have been well studied, albeit mainly independently, the mechanism underlying the linkage of functional dynamics in DMN and MDN with advanced aging is still unclear, as are its effects on cognition across the adult lifespan. One crucial factor affecting functional dynamics is structural integrity of the brain (Warbrick et al., 2017). Accumulating evidence has linked age-related decline of white matter (WM) structural connectivity (fractional anisotropy, FA) to task-related BOLD activity (Brown et al., 2019; Hakun et al., 2015; Webb et al., 2019, 2020; Zhu et al., 2015). In regions of the MDN, some studies have reported that age-related decline in FA is accompanied by “over-recruitment” of BOLD activity in older relative to younger adults during task performance (Burzynska, Garrett, Preuschhof, Nagel, Li, Backman, et al., 2013; Zhu et al., 2015). Longitudinal findings confirm this pattern, in that reduction of FA in the anterior corpus callosum over three years was associated with increased BOLD activity in the ventral lateral prefrontal cortex (Hakun et al., 2015). These results suggest a close association between regions of functional activation and the integrity of the WM fibers connecting them. Likewise, age-related reductions in FA of tracts inter-connecting DMN regions were associated with less DMN deactivation. Yet, most findings demonstrating a relationship between “functional over-recruitment” and WM decline link it to worse behavioral performance (Brown et al., 2015; Brown et al., 2019; Hakun et al., 2015; Zhu et al., 2015). Whether this cross-modality structure-function association is merely a result of structural degeneration or a compensatory mechanism to offset declining cognition is therefore unclear.

Considering that brain activation is more dynamic than brain structure, it is appealing to hypothesize that activation might dynamically change to compensate for degraded brain structure (Dasalaar et al., 2013; Davis et al., 2008). The notion of compensatory activation is not new (Cabeza, 2002; Grady, 2012; Reuter-Lorenz et al., 2000); rather, it is an influential theoretical concept in the cognitive neuroscience of aging, despite arguably insufficient direct empirical validation. To clarify the definition of “compensation” and distinguish it from previous conceptualizations, here we consider “structure-function compensation”, a process that, compared to its absence, results in significantly better cognitive outcomes in older adults. For example, increasing BOLD modulation would be associated with *better* cognitive scores for those with age-related degenerated WM tracts should that modulation be compensatory in nature.

One factor must be considered before further interpretation of the effect of structure-function compensation on cognition: degradation of the structure (e.g., white matter) and the function (e.g., BOLD) of the same brain network does not necessarily occur simultaneously. Notably, structural connectivity between DMN areas evidences age-related declines at a slower (cross-sectional) pace than does structural connectivity in the multiple-demand network (Figley et al., 2015; Hoagey et al., 2019), whereas age-related decline in functional activity occurs earlier in the lifespan in the DMN than the MDN (Park et al., 2010; Park & Friston, 2013; Rieck et al., 2017). These findings are seemingly inconsistent with the previous directional assumption of “*loss of structural integrity, loss of function”* in the DMN (Andrews-Hanna et al., 2019). Therefore, it is worth exploring whether the decline of functional activity in DMN in middle-aged adults is associated with degeneration of WM in the MDN. In other words, exploring the structure-function association between NegMod and FA of MDN-related WM tracts.

To better examine a compensation effect within and across networks, structural and functional data must be acquired separately from each network. While separating BOLD signals according to different brain networks is common, locating white matter tracts that connect the regions belonging to a specific brain network is not. Studies using functionally-guided tractography (which locates WM tracts based on functional regions of interest (ROIs), nodes in functional networks, or clusters demonstrating BOLD activation maxima) have the advantage of focusing on tracts relevant to a “single network,” obviating potential overlap with other networks. Using this method, robust within-network structure-function associations have been reported in DMN and MDN (Brown et al., 2015; Brown et al., 2018; Brown et al., 2019), while studies applying anatomical atlases in tractography have reported inconsistent results (Webb et al., 2019, 2020). The integrity of the tracts connecting functional ROIs of interest might, therefore, be more sensitive to the functional activity adjacent to them (Gazes et al., 2020; Preti et al., 2014; Reid et al., 2017; Staempfli et al., 2008).

Thus far, no study has examined structure-function compensation effects between the DMN and MDN and cognitive function. Therefore, the current study aims to test this account of compensation. We hypothesize that increased greater BOLD signal in response to increases in task difficulty is associated with degraded white matter integrity within the MDN, but is age-dependent (e.g., for middle-aged and older adults) and associated with better cognitive performance. To test these predictions, functionally-guided tractography was employed to locate structural connections between regions showing age-related BOLD modulation (see Rieck et al., 2017). We tested our hypotheses by (1) locating WM tracts connecting to a major hub in each network (insula for PosMod and precuneus for NegMod) and examining the tracts for age-related differences in microstructure; (2) for the tracts identified, testing whether there are age-related structure-function associations within and across the two networks; (3) examining whether and how age-related structure-function associations are associated with a measure of executive function. Three possible associations would qualify as a compensatory effect: *within-network across modality compensation* (MDN-sourced BOLD and FA; DMN BOLD and FA), *between-network across modality compensation* (MDN BOLD and DMN FA; DMN BOLD and MDN FA), and/or *coupled BOLD network modulation and tract FA from either network* (MDN x DMN BOLD and FA from MDN or DMN tracts).

The present study aims to test the cross-network structure-function associations and the dynamic alteration of BOLD modulation between the MDN and DMN in offsetting degenerated WM microstructure in the MDN. We hypothesized that WM tracts in the MDN would show significant negative age effects and, according to current evidence on structure, this degraded WM would be associated with weakened BOLD modulation in DMN for middle-aged and older adults. Conceptually (see Figure 1), previous findings support a “non-compensatory” effect of functional activity on cognition, where age-related decreases in BOLD modulation to task are associated with worse cognition in older age (Kennedy et al., 2017; Rieck et al., 2017). The current study posits a “compensatory” effect of functional activity coupled between the two networks in response to degenerated WM in MDN. Specifically, BOLD modulation of the MDN is expected to increase, to be coupled with decreased BOLD modulation of the DMN, and to predict better cognitive performance in middle-aged and older adults.

**Figure 1.**
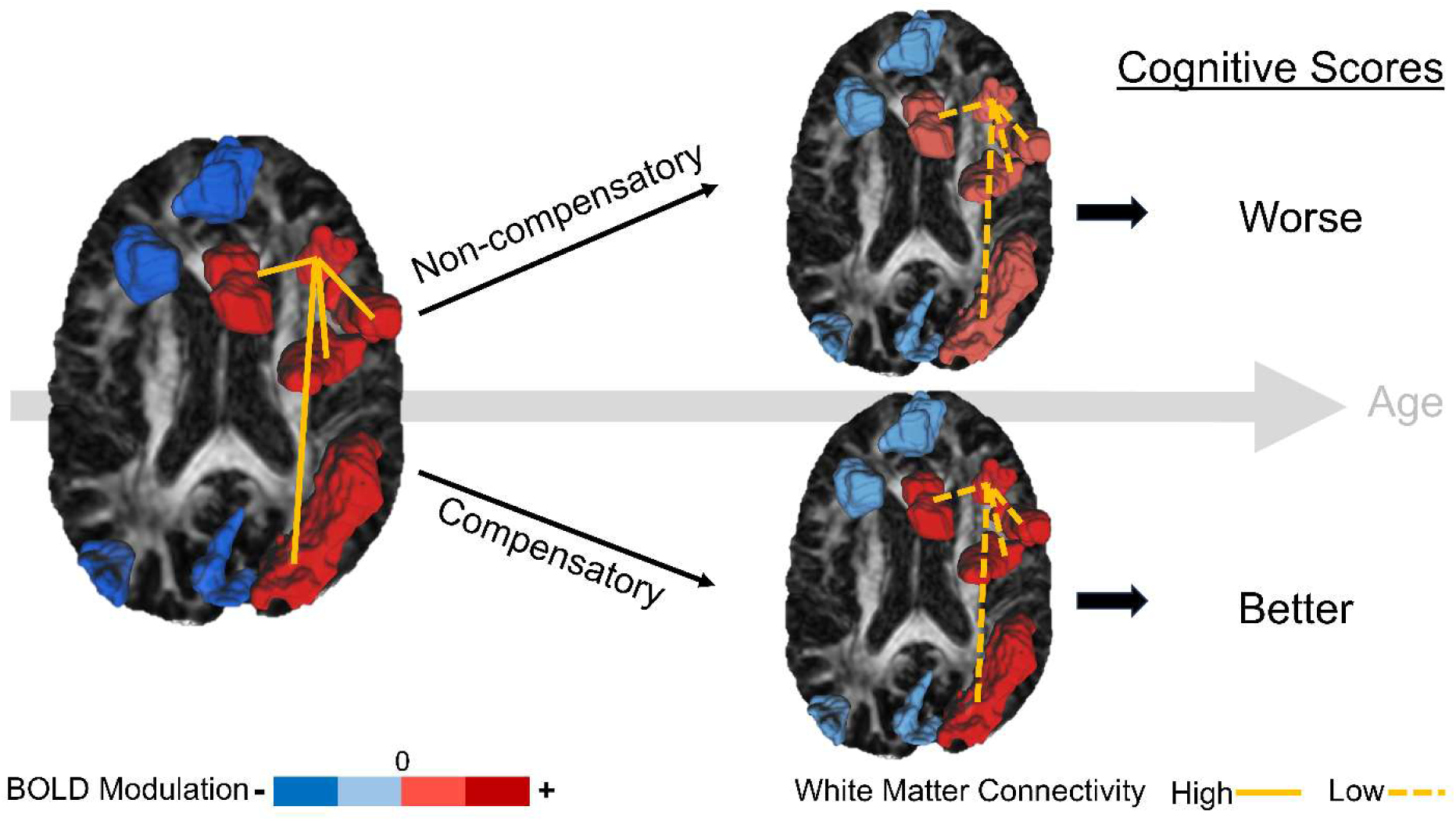
Conceptual hypothesis of structure-function association compensatory effect on cognitive performance. The nine ROIs in each brain are the functional clusters showing the effect of age-related BOLD modulation on task difficulty; the BOLD modulation legend illustrates the direction and strength of the effect where dark red/blue color indicates strengthened positive/negative BOLD modulation, respectively, while the lighter color represents weakened BOLD modulation effect as a result of brain aging. The four solid/dashed lines in each brain represent relative intact/degraded white matter connections from the hub of the multiple demand network (MDN), the insula, to the other four clusters in the MDN. The brain on the left represents the state of a healthy, younger brain, with relatively high white matter connectivity and high positive/negative BOLD modulation to task difficulty, both of which are linked to better performance. As the brain ages, white matter in the MDN degrades first (solid lines connecting red ROIs in younger adults become dashed lines with aging, on the brains on the right side). In response to this structural impairment in frontal-parietal regions, the functional activity could demonstrate either a “Non-compensatory” relation to cognition, where declines in BOLD modulation in both networks are associated with worse cognition, as the top right brain shows, or a “Compensatory” effect, where strengthened local activity (increased PosMod) coupled with weakened NegMod is linked to better cognition (bottom right brain).

## 2. METHODS

### 2.1. Participants

This cross-sectional study utilized a lifespan sample of 161 healthy adults from the first wave of our Dallas Area Lifespan Longitudinal Aging Study (D.A.L.L.A.S), the same sample of participants as described in Rieck et al. (2017). Participant ages ranged from 20-94 years (mean age = 51.93 ± 18.9 yrs; 59% women) and they were recruited from the Dallas-Fort Worth metroplex. Participants were screened to be right-handed with normal or corrected-to-normal vision and hearing. To screen against signs of dementia and depression, all participants underwent a Mini Mental Status Exam (MMSE) (Folstein et al., 1975) and obtained a score of at least 26 (mean MMSE = 29, range 26 - 30); they also completed the Center for Epidemiologic Study Depression Scale (CESD; Radloff (1977)) and were required to score less than or equal to 16 on the scale (mean CESD = 4.29, range 0 - 16). Participants were further screened against a history of neurological, cardiac, metabolic or psychiatric conditions, head injury with loss of consciousness > 10 min, or current psychotropic medication use. Participants completed two visits for neuropsychological and cognitive tests at the University of Texas at Dallas’ (UTD) Center for Vital Longevity and an MRI session at the Advanced Imaging Research Center at the University of Texas Southwestern Medical Center (UTSW). The study was approved by the UTD and the UTSW Institutional Review Boards. Participants provided written informed consent before entering the study and they were compensated for their time. See **Table 1** for participant demographics.

**Table 1.** Participant demographics.

|  | Younger | Middle-Aged | Older | Oldest | Whole Sample |
| --- | --- | --- | --- | --- | --- |
| Age range | 20–35 | 36–54 | 55–69 | 70–94 | 20-94 |
| N (% female) | 42 (54.8%) | 41 (56.1%) | 40 (62.5%) | 38 (63.2%) | 161 (59.0%) |
| Age Mean (± sd) | 27.45 (4.4) | 45.24 (5.4) | 61.05 (3.7) | 76.58 (6.1) | 51.93 (18.92) |
| Education Mean (± sd) | 15.57 (2.1) | 15.12 (2.5) | 15.65 (2.4) | 15.71 (2.7) | 15.51 (2.4) |
| MMSE Mean (± sd) | 29.14 (.95) | 29.32 (.79) | 28.95 (.72) | 28.87 (.84) | 29.07 (.84) |
| CESD Mean (± sd) | 4.60 (3.66) | 4.80 (4.13) | 4.20 (3.64) | 3.50 (3.69) | 4.29 (3.78) |

*Note.* Participants were separated into age groups for illustration purposes; age was treated as continuous variable in all statistical models. sd: standard deviation; MMSE: Mini Mental Status Examination; CESD: Center for Epidemiologic Study Depression Scale.

### 2.2. Cognitive Measures

For the current study, three metrics of cognitive performance were selected as outcome measures. From the fMRI task (see section 2.3.2), mean accuracy of the distance judgments (averaged across conditions and across runs) was computed as an index of in-scanner task performance. As measures of executive function, switching and inhibition composites were computed from two subtests of the Delis-Kaplan Executive Function System (Delis et al., 2001) (D-KEFS): Verbal Fluency and Stroop Color-Word Interference tasks. Scores from the “Culture Fair Intelligence Test” (2007) (CFIT) served as an index of fluid intelligence. Total time taken for each task was about 10 – 12 minutes. Individual cognitive test details are described below.

#### 2.2.1. D-KEFS Verbal Fluency Category Switching task

This task contained three conditions: Letter Fluency, where participants generated as many words as possible in 60 seconds that begin with a specific letter; Category Fluency, which required participants to generate as many words as possible in 60 seconds belonging to a specified category (e.g. Animals). To suppress retrieving repeated and/or irrelevant responses, inhibition may be involved in these two conditions (Shao et al., 2014); and Category Switching, which required participants to switch back and forth between naming words belonging to one of two different specified categories (e.g., apple, chair, banana, desk, …etc) for 60 seconds. As an index of Inhibition, the mean number of correct responses for letter fluency and category fluency (3 letter trials, 2 category trials) was calculated. The index of Switching was the total correct score for the Category Switching condition.

#### 2.2.2. D-KEFS Color-Word Interference (Stroop) task

The Stroop task contained four conditions: Two control conditions included Color Naming speed and Word Reading speed, where participants either named the color of the patches or read the words printed in black. In the Inhibition condition, they were asked to name the color of the ink, regardless of the actual word. In the Inhibition/Switching condition, participants had to switch between naming the ink color or reading the word. Response time (seconds) to complete was recorded in each condition. The control condition score was calculated as the mean completion time of color and word naming. The “inhibition score” or “switching score” for this task was calculated by subtracting the control condition score from either Inhibition or Inhibition/Switching condition RT. All three scores were multiplied by −1 before further processing.

To create general composite scores of switching and inhibition performance, the switching and inhibition condition scores from the two tasks were first transformed to z scores, and then averaged across-task, within-domain (i.e., switching composite = [z fluency switching score + z stroop switching score] / 2).

#### 2.2.3. Fluid Reasoning: CFIT

The Cattell Culture Fair Intelligence Test (CFIT) is designed to evaluate fluid intelligence, particularly reasoning. The test utilizes matrix reasoning problems and consists of four subtests each targeting different aspects of fluid intelligence. Scale 3 and Form B were used in the current study, containing a total of 50 items across four subtests: 1) Series subtest (13 items) assesses the ability to recognize and extend patterns in a sequential manner; 2) Classification subtest (14 items) measures individual’s capacity to discern subtle differences and categorize objects. 3) Matrices subtest (13 items) evaluates the ability to identify missing elements; 4) Topology subtest (10 items) measures participants’ analogous and spatial abilities. Correct responses within the time limit were recorded across the four subtests and averaged to create a composite fluid intelligence score (CFIT score).

### 2.3 MRI Protocol

#### 2.3.1. Sequence Acquisition

All participants were scanned on a single Philips Achieva 3T scanner equipped with a 32-channel SENSE head coil. BOLD data were collected using a T2*-weighted echo-planar imaging sequence (EPI) with 29 interleaved axial slices parallel to AC-PC line. Matrix size = 64 × 64 × 29, voxel size = 3.4 × 3.4 × 5 mm^3^, field of view (FOV) = 220 mm, TE = 30 ms, and TR = 1500 ms. High resolution anatomical T1-weighted images were collected with an MP-RAGE sequence with the following parameters: 1 × 1 × 1 mm^3^, FOV = 256 mm, TE = 3.8 ms, TR = 8.3 ms, flip angle = 12°. DTI data were collected using a single shot EPI sequence with 65 axial slices, voxel size 2 × 2 × 2.2 mm^3^ (reconstructed to 0.88 × 0.88 × 2.2 mm^3^), 30 diffusion-weighted directions, b-value = 1000 s/mm^2^, 1 non-diffusion weighted image (b = 0 s/mm^2^); TR/TE = 5608/51 ms, FOV = 224 ×224 ×143 mm^3^, matrix = 112 × 112 mm^2^.

#### 2.3.2. fMRI distance judgment task procedure

Participants performed in the scanner a spatial distance judgment task (Baciu et al., 1999; Park et al., 2010). In a blocked design, participants performed two types of judgments: (1) in the control condition, a horizontal bar appeared in the middle of the screen, accompanied with a dot on either the left or right side of the bar, and participants indicated whether the dot was on the left or right with a button press; (2) At the beginning of each task block, participants were cued with a vertical bar in the middle of the screen, which served as a reference of distance, for three seconds, then disappeared and was replaced by a horizontal bar and a dot either above or below it. Their task is to judge whether the dot was “nearer” or “farther” from the bar relative to the reference bar length. This judgment involved three levels of difficulty: easy, medium and hard, parametrically varying the distance between the dot and bar. The reference cue was only displayed before each block of trials and did not vary. Participants responded with their index or middle finger whether the dot was “farther”/ “nearer” or “left”/ “right” relative to the bar. Stimulus presentation time was 2500 ms with a 500 ms interstimulus interval. There were three runs with 20 blocks in each run, and four conditions were presented in a pseudo-random order in each run. The total functional scan time was about 15 min. Mean accuracy and median response time (RT) for each task condition (easy, medium, hard) were reported in our previous publication (Rieck et al., 2017). For the present study, mean accuracy across task difficulty (easy, medium, hard conditions) was the index of performance.

##### 2.3.2.1. fMRI analysis

The current structure-function study used the previously processed, analyzed, and published functional clusters from the age-related BOLD modulation to difficulty effect reported in Rieck et al., (2017). Detailed image processing steps are included in our previous study and here we only include details pertinent to the current study. Increased and decreased activation in the Hard relative to the Easy distance judgment conditions was the contrast of interest. That contrast isolated voxels that increased or decreased their activation with increasing task difficulty, a process henceforth referred to as “BOLD modulation”. Increased activation to greater difficulty is referred throughout as “Positive BOLD Modulation”; decreased activation to greater difficulty is referred to as “Negative BOLD Modulation”. fMRI data were all transformed to MNI space (unlike the diffusion data described below, which were kept in native space). The second-level model included age as a mean-centered continuous predictor to isolate age effects on positive and negative BOLD modulation in a whole brain voxel-wise manner with a conservative height threshold p < .0001, and clusters accepted as reliable at an extent threshold of p < .05 FWE. Those resulting clusters that demonstrated reliable age differences on BOLD modulation to difficulty were taken directly from (Rieck et al., 2017) and re-shown here in **Table 2**. There were 5 clusters where activity increased with greater difficulty (“positive modulation”, all were MDN regions) and 4 clusters that decreased in activation with greater difficulty (“negative modulation”, all were DMN regions). Perusal of **Table 2** suggests a lateralization such that the positive modulation (MDN) regions were right-lateralized while the negative modulation (DMN) regions were mostly left-lateralized or on the midline. Thus, these within-network clusters are also mainly within-hemisphere, whereas interactions between-networks to be tested in this study are cross-hemispheric. Mean BOLD parameter estimates were extracted from each cluster listed in **Table 2** using the MarsBaR toolkit (Brett et al., 2002) and used for all subsequent analyses.

**Table 2.** Cluster peaks for the regression of age on modulation of BOLD activation to difficulty (from Rieck et al., 2017)

| ROI# | Cluster label | BA | *k* | *x* | *y* | *z* | *t* | *p*<sub>FWE</sub> |
| --- | --- | --- | --- | --- | --- | --- | --- | --- |
| **Positive modulation (hard > easy)** |  |  |  |  |  |  |  |  |
| 1 | R intraparietal sulcus, inferior & superior parietal lobules | 40 | 603 | 45 | -39 | 54 | 5.70 | < .001 |
| 2 | R insula | 13 | 176 | 42 | 21 | -6 | 5.35 | < .001 |
| 3 | R middle frontal | 6 | 86 | 33 | 0 | 57 | 5.13 | 0.003 |
| 4 | R inferior frontal | 44 | 94 | 51 | 9 | 24 | 5.03 | 0.002 |
| 5 | R anterior cingulate | 32 | 65 | 6 | 18 | 48 | 4.64 | 0.007 |
| **Negative modulation (easy > hard)** |  |  |  |  |  |  |  |  |
| 6 | L middle frontal | 9 | 69 | -27 | 33 | 42 | 5.41 | 0.006 |
| 7 | L angular gyrus | 39 | 91 | -45 | -75 | 33 | 5.35 | 0.002 |
| 8 | L/R cuneus, precuneus & posterior cingulate | 18 | 241 | -3 | -87 | 24 | 5.20 | <.001 |
| 9 | L/R ventromedial prefrontal | 10 | 135 | 0 | 51 | -6 | 4.76 | <.001 |

*Note.* Height threshold of $p < .0001$ and cluster correction at $p < .05$ , family-wise error corrected. BA – Brodmann area; *k* - # voxels in cluster; R - Right; L - left

#### 2.3.3 DTI Preprocessing and Analysis

DTI data were preprocessed using the DTIPrep v1.2.4 quality control package. Diffusion tensors were computed using DSI Studio (Yeh et al., 2013). The nine grey matter functional clusters in **Table 2** served as our functionally-informed ROIs for deterministic tractography (implemented in DSI Studio) to build WM tracts of interest traversing among these functional regions. To allow tractography to be performed in native diffusion space the fMRI clusters were non-linearly registered to each participant’s b0 diffusion image with the Advanced Normalization Tools package (Avants et al., 2009) using a registration of the MNI 1mm^3^ template to each participant’s T1-w MPRAGE, followed by a non-linear registration between the T1-w MPRAGE and the b0 diffusion image using linear interpolation. Each functional cluster was warped from MNI template space to the participant’s diffusion space simultaneously using the linear and non-linear warps with nearest neighbor interpolation. These warped ROIs were then dilated to aid in tractography. The mean number of white matter voxels within these raw clusters was 1284 voxels. Each ROI was then dilated with a kernel size of 0–3, so that they all covered at least 1284 white matter voxels.

##### 2.3.3.1. Deterministic tractography

To build white matter connections that were the most related to the positive BOLD modulation in MDN and negative BOLD modulation in DMN, we focused on the connections linked to the hub of the respective network: either the insula (cluster #2 from **Table 2**) or cuneus/precuneus (cluster # 8 from **Table 2**) for MDN and DMN, respectively (Bullmore & Sporns, 2009). All 15 possible cluster combinations were submitted to fiber-tracking in each participant. Deterministic tractography was performed for each participant; each tract generated was carefully visually inspected, and any grossly atypically appearing tracts were excluded from analysis. The DSI Studio tractography fitting parameters for each tract can be found in **Supplemental Table 1**. To ensure a multivariate normal distribution of number of streamlines among all tracts, the Mahalanobis distance was calculated using the number of streamlines for each tract for each participant using R v4.1.0. Participants with extreme Mahalanobis distance on their streamlines (using *pchisq* in R, p < .001), indicating that the number of streamlines of one or more tracts was either extremely low or high, were categorized as outliers and removed from further analysis. In total, nine participants were removed for low streamline counts.

###### Tract Visualization and Identification

Of the possible 15 combinations of functional ROI pairs, 5 robust functionally-guided tracts were identified and are illustrated in **Figure 2**. Two of these tracts resided within the PosMod-related network, identified in red in the figure. The first tract connected right posterior parietal cortex with right anterior insula (i.e., ROI #s 1 and 2 in Table 2), and because it most closely overlaps anatomically with the inferior frontal occipital fasciculus (IFOF), which is a deep white matter long-range association tract, we refer to it as “IFOF” in the current study. The second within-PosMod tract identified was a small U-fiber tract (which is a short-range superficial white matter tract) connecting the right anterior insula with the right inferior frontal gyrus (IFG) pars opercularis (i.e., ROI #s 2 and 4), which we named fronto-insular “U-fiber” (Catani & Thiebaut de Schotten, 2008).

**Figure 2.**
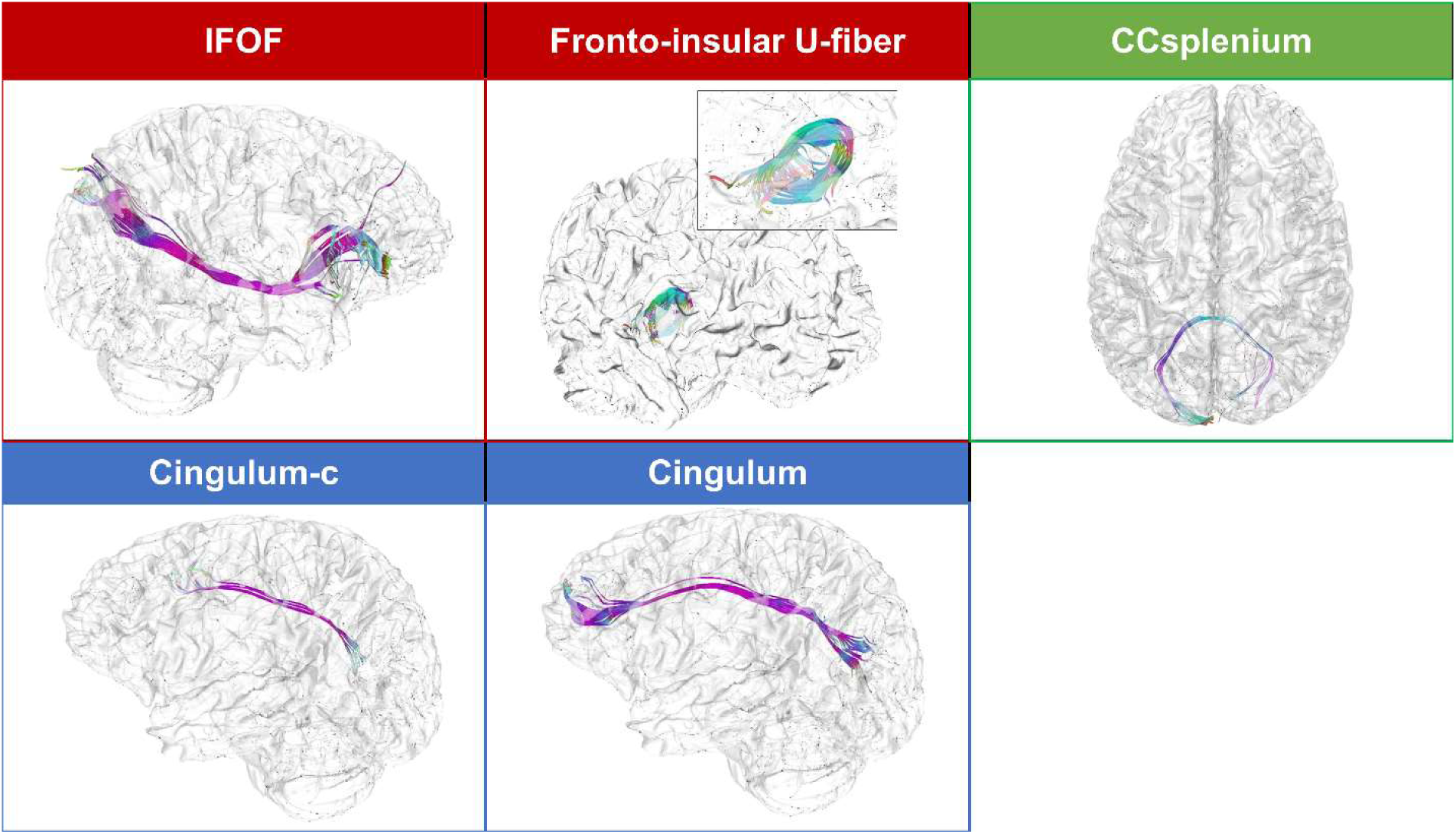
Overview of tracts obtained using functionally-defined ROIs. Tracts resulting from functionally-guided tractography were named by the anatomical tract with which they most overlapped. Tracts residing between functional clusters within the positive network are boxed in red, tracts between negative network clusters are boxed in blue, and the tract resulting from cross-hemispheric connection between networks is boxed in green. All tracts are projected on the isosurface of the brain for presentation purposes, which is registered using participant’s T1-weighted image in DSI-studio. CC – corpus callosum, Cingulum-c – tract from between network, cross-modulation.

For the within-NegMod-related network one tract was identified (blue box in **Figure 2**), connecting left midline/bilateral cuneus/precuneus with left midline/bilateral ventromedial prefrontal cortices (i.e., ROIs # 8 and 9). This tract overlaps most closely anatomically with the long-range association cingulum bundle and was named “Cingulum”.

Finally, two of the generated tracts interconnected a PosMod region to a NegMod region, i.e., between-network (green box in **Figure 2**). The first tract connected right posterior parietal cortex with left precuneus/posterior cingulate (thus traversing the corpus callosum, CC) and largely overlapped with CC splenium inter-hemispheric fibers, hence it was named “CCsplenium”. The second tract was largely redundant with the “Cingulum” tract described above, as it connected the right anterior cingulate and the left precuneus/posterior cingulate (ROIs #5 and 8) and is labelled “Cingulum-c” (cross-networks) in **Figure 2**. However, due to the anatomical overlap between the “Cingulum-c” and “Cingulum”, the shorter Cingulum-c tract was excluded from further analysis.

To statistically control for the effects of global white matter FA, a mean whole brain FA (WBFA) metric was additionally created for each participant as follows. Whole brain white matter streamlines were generated using DSI Studio with the following tractography fitting parameters: no regional restrictions were applied, tract lengths between 20 to 500 mm, turning angle = 60; all other parameters were set as specified for the previous tracts. From the resulting whole brain streamlines, FA was extracted and averaged across all voxels. This WBFA metric was entered into statistical models as a covariate of no interest.

### 2.4. Statistical approach

To test the three study aims (detailed in each corresponding results section) General Linear Models (GLM) were specified and computed using R v4.1.0 (R core team, 2021). Quadratic age and FA terms were examined based on previous studies that reported quadratic effects of age on white matter microstructure (Slater et al., 2019). WBFA served as a covariate in all models. The two BOLD modulation variables were the mean of the contrast parameter estimates of the five PosMod-related clusters or the mean estimates of the four NegMod-related clusters. To examine the structure-function compensation effects on executive function and fluid intelligence (aim 3), we ran separate models employing age, FA, FA^2^, positive/negative BOLD modulations, and their interactions as predictors of task accuracy, executive function, or fluid intelligence as the three dependent variables, respectively.

Significant interaction terms involving continuous variables were decomposed using simple slopes and the Johnson-Neyman procedure (Neyman & Pearson, 1936). False discovery rate (FDR) was used to correct for multiple comparisons of each common term across models (e.g., p-values of the main effects of age of all models were grouped together and corrected) using the R function “p.adjust” (Benjamini & Hochberg, 1995). All significant effects described in the results section survived correction unless otherwise specified, and all p values are given prior to correction.

## 3. RESULTS

### 3.1. Age Effects on White Matter Tract FA, Controlling for Whole Brain FA

To examine the cross-sectional age effect on WM microstructure within/between networks, GLMs were specified with each tract FA as its dependent variable, linear and quadratic age as the simultaneous continuous predictors, and WBFA^1^ as covariate of no interest (formula a).

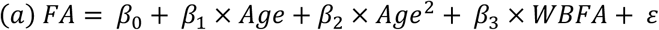

The frontal-insular U-fiber tract uniquely demonstrated a significant negative linear age effect (t(128) = −3.65, p <.001, η^2^ = 0.09), indicating that increasing age was associated with decreasing FA. The other three tracts did not evidence significant age effects after controlling for WBFA (**Figure 3**). WBFA was significantly associated with tract FA (after accounting for age) for all but CCsplenium. However, the linear age effect on FA of the four tracts were all significant before controlling for WBFA (t(113) = −2.7, p < .001; t(129) = −5.87, p < .001; t(101) = −2.76, p < .001; and t(94) = −4.07, p < .001, for IFOF, U-fiber, CCsplenium, and Cingulum, respectively). No significant quadratic age effects were observed for any tract. See **Table 3** for summary statistics for the four models and **Figure 3** for age-FA scatterplots (adjusted for WBFA).

**Figure 3.**
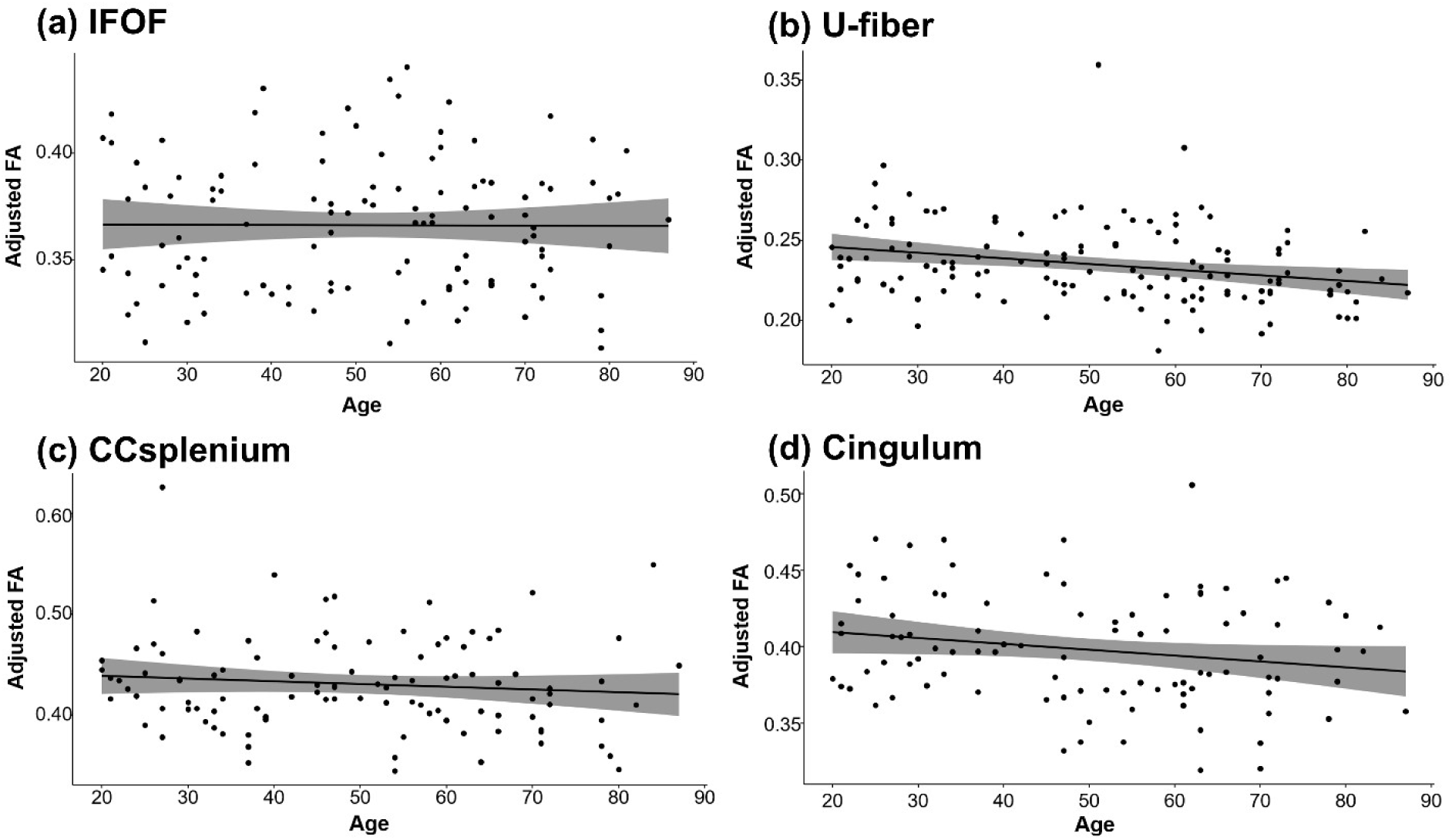
Correlation of Age and FA after controlling for whole brain FA. Significant age-related decline of FA after controlling for whole brain FA in the frontal-insular U-fiber tract is observed, but not for the remaining three functionally-guided tracts: the inferior frontal-occipital fasciculus, the splenium of the corpus callosum, and the cingulum bundle (see also Table 3).

**Table 3.**
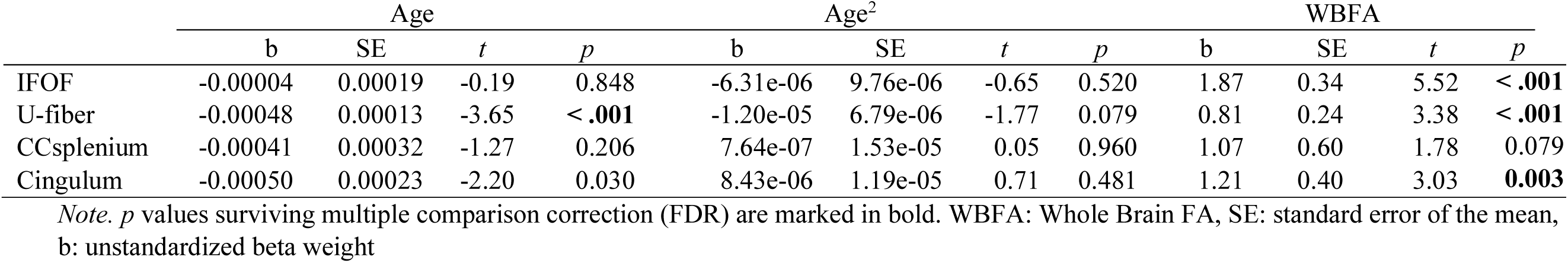
Summary of effects in the regression models. Fractional anisotropy (FA) is estimated with Age, quadratic effect of Age, and Whole brain FA for each functionally-guided white matter tract.

|  | Age |  |  |  | Age<sup>2</sup> |  |  |  | WBFA |  |  |  |
| --- | --- | --- | --- | --- | --- | --- | --- | --- | --- | --- | --- | --- |
|  | b | SE | *t* | *p* | b | SE | *t* | *p* | b | SE | *t* | *p* |
| IFOF | -0.00004 | 0.00019 | -0.19 | 0.848 | -6.31e-06 | 9.76e-06 | -0.65 | 0.520 | 1.87 | 0.34 | 5.52 | < .001 |
| U-fiber | -0.00048 | 0.00013 | -3.65 | < .001 | -1.20e-05 | 6.79e-06 | -1.77 | 0.079 | 0.81 | 0.24 | 3.38 | < .001 |
| CCsplenium | -0.00041 | 0.00032 | -1.27 | 0.206 | 7.64e-07 | 1.53e-05 | 0.05 | 0.960 | 1.07 | 0.60 | 1.78 | 0.079 |
| Cingulum | -0.00050 | 0.00023 | -2.20 | 0.030 | 8.43e-06 | 1.19e-05 | 0.71 | 0.481 | 1.21 | 0.40 | 3.03 | **0.003** |

Note. *p* values surviving multiple comparison correction (FDR) are marked in bold. WBFA: Whole Brain FA, SE: standard error of the mean, b: unstandardized beta weight

### 3.2. Brain Structure-Function Association: Relation between Age-related White Matter Tract FA and BOLD Modulation to Difficulty

To examine structure-function associations, i.e., how WM microstructure relates to BOLD modulation, GLMs were specified with positive or negative BOLD modulation as the dependent variable, and age, FA, FA^2^, and WBFA entered as predictors as well as the interaction of age × FA and age × FA^2^. Formula b provides an example of a model for positive BOLD modulation. Separate models were constructed for each of the 4 WM tracts, for each of the 2 types of BOLD modulation (negative and positive) for a total of 8 models.

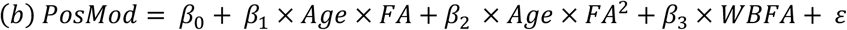

#### Positive BOLD Modulation and White Matter

As reported in Rieck et al., 2017, there was a negative effect of age on positive BOLD modulation (all p’s < .01). None of the four WM tract FA terms was significantly related to positive BOLD modulation after FDR correction, although there were trends for a quadratic effect of IFOF FA on PosMod and the age x IFOF FA interaction. See Supplemental Table 2a for detailed model statistics. These results suggest that the microstructural organization of these functionally-informed WM tracts may not play a significant role in task-positive network modulation to difficulty.

#### Negative BOLD Modulation and White Matter

Again, there was a detrimental effect of age on negative BOLD modulation (reported in Rieck et al., 2017, and see **Supplemental Table 2b** for all model statistics). Of the four models, there was one tract, IFOF, that evidenced a significant FA-BOLD association, and this association was age dependent (age x IFOF FA interaction: *b* = 0.28, *t*(120) = 3.32, *p* = .001, η^2^ = 0.08). None of the other tracts showed FA-NegMod associations. To determine the nature of the IFOF FA x age interaction, simple slopes and the Johnson-Neyman procedure were used (see **Figure 4**). For younger individuals (1 SD below the mean age), FA of the IFOF tract was *inversely* related to NegMod, where higher FA was associated with weaker negative modulation to difficulty (and less so as FA increased). For middle-aged adults (at the mean age), IFOF FA was not associated with BOLD modulation. Strikingly, for older individuals (1 SD above the mean age), a *positive* association was observed such that lower IFOF FA was associated with relatively stronger negative modulation to difficulty (**Figure 4a**). This moderation effect is evident for younger adults up until approximately age 34, and for older individuals beginning around age 62. In other words, older adults who demonstrated degraded white matter in a tract interconnecting PosMod-related activation clusters (IFOF) demonstrated strengthened negative modulation, whereas there was no such association for middle-aged adults, and the opposite pattern was seen in younger adults.

**Figure 4.**
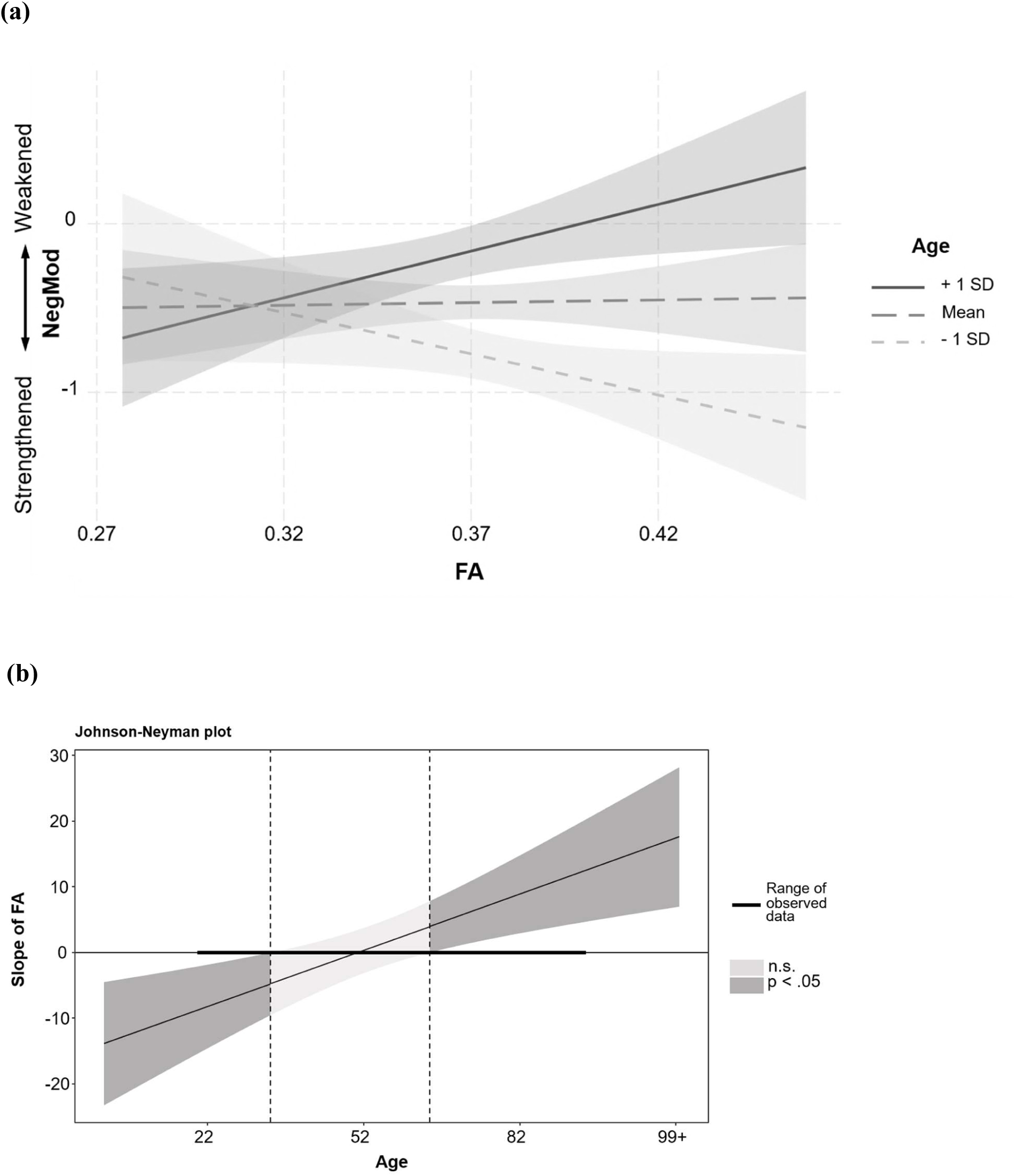
The nature of the white matter structure – BOLD function association depends upon age. Breakdown of the significant Age x IFOF FA interaction on negative BOLD modulation to difficulty. (a) Simple slopes plot illustrating that the BOLD-FA association is positive in older adults (1 SD > mean age), with higher anisotropy associated with higher negative BOLD modulation (a weakened dampening of cortical activation); nonsignificant for middle-aged adults (mean age), and negative for younger adults (1 SD < mean age), with higher anisotropy associated with weaker negative BOLD modulation (stronger dampening of cortical activation) to difficulty. (b) Johnson-Neyman plot indicates that this association is significant in younger adults until ∼age 34 and in individuals above ∼age 64.

### 3.3. Effects of Age, WM Tract FA, and BOLD Modulation on Cognitive Performance

The third, and final goal of the study was to examine whether age-related structure-function effects were positively associated with cognitive performance, i.e., whether there is support for structure-function compensation, a compensatory role of functional activity with white matter degradation. Models were tested with either fMRI task accuracy, switching, inhibition, or fluid intelligence as the dependent variable, and with age, positive/negative BOLD modulation, and FA/quadratic FA and their interactions as independent variables, while controlling for WBFA (see formula c for an example of model predicting CFIT performance).

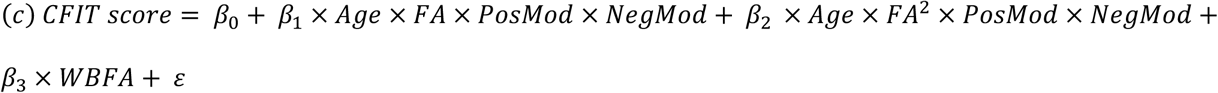

Regression models were specified for each of the four WM tracts for each of the four cognitive scores, for a total of 16 models. Detailed model result parameters/statistics are provided in **Supplemental Table 3a-3d**, and a conceptual summary of all significant results is presented in **Table 4**.

**Table 4.**
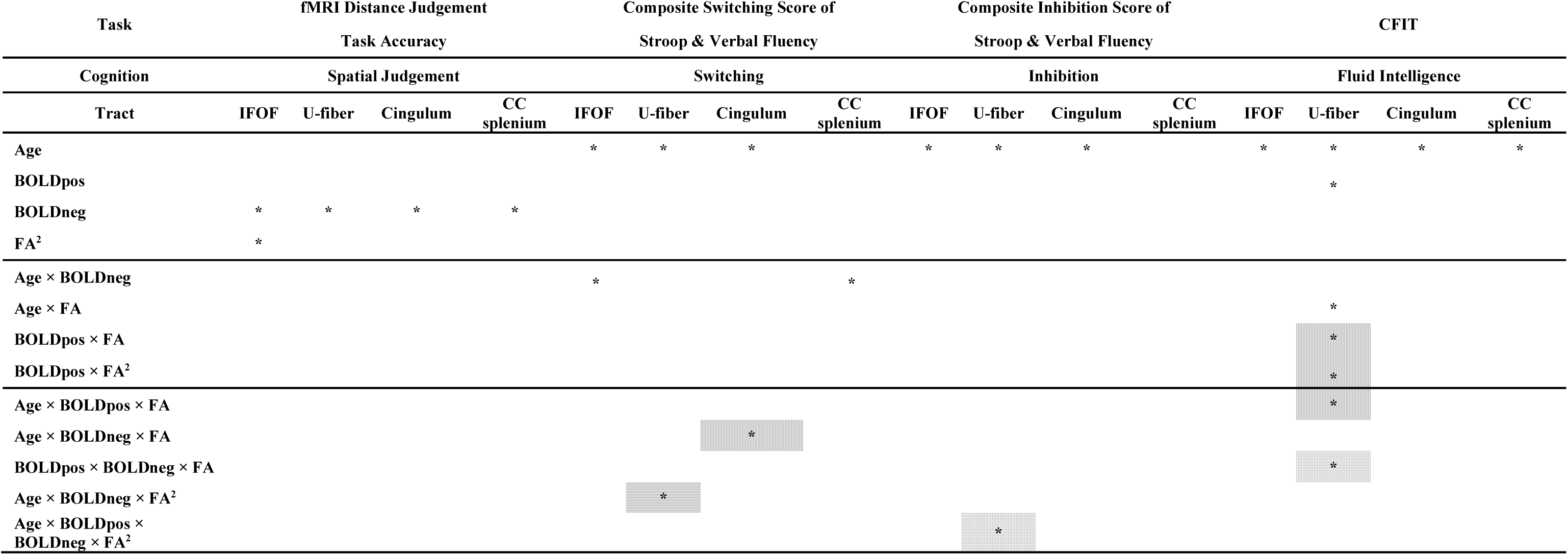
Summary of significant results: Effects of Age, PosMod, NegMod, linear and quadratic FA and their interactions on four cognitive constructs after controlling for WBFA. Significant results surviving multiple comparison correction using FDR are labelled with asterisks. Cells marked with vertical stripes are interaction effects representing “*within-network structure-function compensation*”, where both FA and BOLD were retrieved/generated from the same functional network; cells filled with horizontal stripes represent “*cross-network structure-function compensation*”, where FA and BOLD were retrieved from different functional networks; grid patterns show “*coupling effects”*, where FA modulates both BOLDpos and BOLDneg on cognitive scores. BOLDpos: positive BOLD modulation to task difficulty, or PosMod; BOLDneg: negative BOLD modulation to task difficulty, or NegMod. Detailed model parameters of the results can be found in supplemental Table 3a-3d.

#### In-scanner Task Accuracy

In the models predicting accuracy on the distance judgment task, equivalent performance across the lifespan was observed (*p* > .01). Whereas magnitude of positive BOLD modulation was unrelated to accuracy, amount of negative modulation was significantly related to accuracy across all four models (*p*’s < .03). Additionally, an effect of quadratic FA on task accuracy was found for the IFOF model (IFOF FA^2^; b = 15.47, t(102) = 2.83, *p* < .006, η^2^ = 0.06; see Supplemental Figure 1). This finding suggests that while FA has a positive effect on accuracy at higher levels of FA, the effect diminished at low to moderate levels of FA and became a negative effect at the lowest level of FA. This effect at lowest levels of FA might be due to a compensatory role of BOLD (low FA-high accuracy in high BOLD individuals), although IFOF FA’s interaction with PosMod and NegMod became non-significant after multiple comparisons correction, see Supplemental Table 3a. The same interactions (in these cases remaining significant after correction) were also identified for out-of-scanner cognitive performance, as detailed in the following section.

#### Out-of-scanner Cognitive Performance

To streamline the presentation of the results, we report findings across cognitive and WM tract models by main effect terms, and then by lower-order interaction terms, followed by the structure-function interaction terms (i.e., the hypothesized compensatory effects).

#### Age main effects

As would be expected, significant main effects of age were observed across all three out-of-scanner cognitive tests, with increasing age associated with decreased performance in switching, inhibition, and fluid intelligence (all *p*’s < .02).

#### BOLD modulation effects

There was a significant main effect of *positive* BOLD modulation on CFIT for the model testing the U-fiber frontal-insular tract (b = 0.62, t(116) = 2.81, *p* = .006, η^2^ = 0.06), indicating that lower positive modulation to task difficulty was associated with poorer CFIT performance. For *negative* BOLD modulation to task difficulty, age × negative BOLD modulation interactions on switching performance were significant for the IFOF (b = −0.02, t(102) = −2.37, *p* = .020, η^2^ = 0.04) and CCsplenium models (b = −0.02, t(88) = −2.46, *p* = .016, η^2^ = 0.05). For the IFOF model, weaker negative BOLD modulation was associated with poorer switching performance for individuals above ∼59 years of age (via Johnson-Neyman interval analysis). For the CCsplenium model, older adults also showed weaker negative BOLD and poorer switching performance. In contrast, younger adults showed the opposite trend, with strengthened negative BOLD modulation associated with poorer switching performance.

#### Structure × Function interaction effects

The structure-function association results detailed below, and summarized in Table 4, can be further categorized into three types of structure-function effects: 1) when FA moderates the BOLD modulation in the same network (*within-network effect*; e.g., MDN tract and MDN BOLD), 2) in a different network (*between-network effect*; e.g., MDN tract and DMN BOLD), 3) or when there is a coupling of both positive and negative BOLD modulation (*coupling effect*; PosMod × NegMod interaction) on cognitive scores.

Within MDN. For *within-network structure-function effect* in MDN, there was a significant interaction of PosMod × U-fiber tract FA on CFIT (b = −25.69, t(116) = −2.59, *p* = .011, η^2^ = 0.05), as well as a PosMod × U-fiber tract FA^2^ interaction (b = −80.22, t(117) = −2.73, *p* = .01, η^2^ = 0.05; see **Figure 5a**). These interactions were qualified, however, by a significant age × PosMod × U-fiber FA interaction (b = −1.47, t(116) = −2.61, *p* = .01, η^2^ = 0.05), which suggests that the structure-function association was age-dependent. The decomposition of this interaction revealed a cross-modality interaction (**Figure 5b**), where age and FA modulate the effect of PosMod on the CFIT score. A positive effect was observed for middle-aged participants with lower levels of FA, where greater positive BOLD modulation was associated with better reasoning performance. However, an adverse effect was observed for middle-aged adults with higher levels of FA. Strengthening positive BOLD modulation was associated with worse performance for participants maintaining relatively high white matter connectivity.

**Figure 5.**
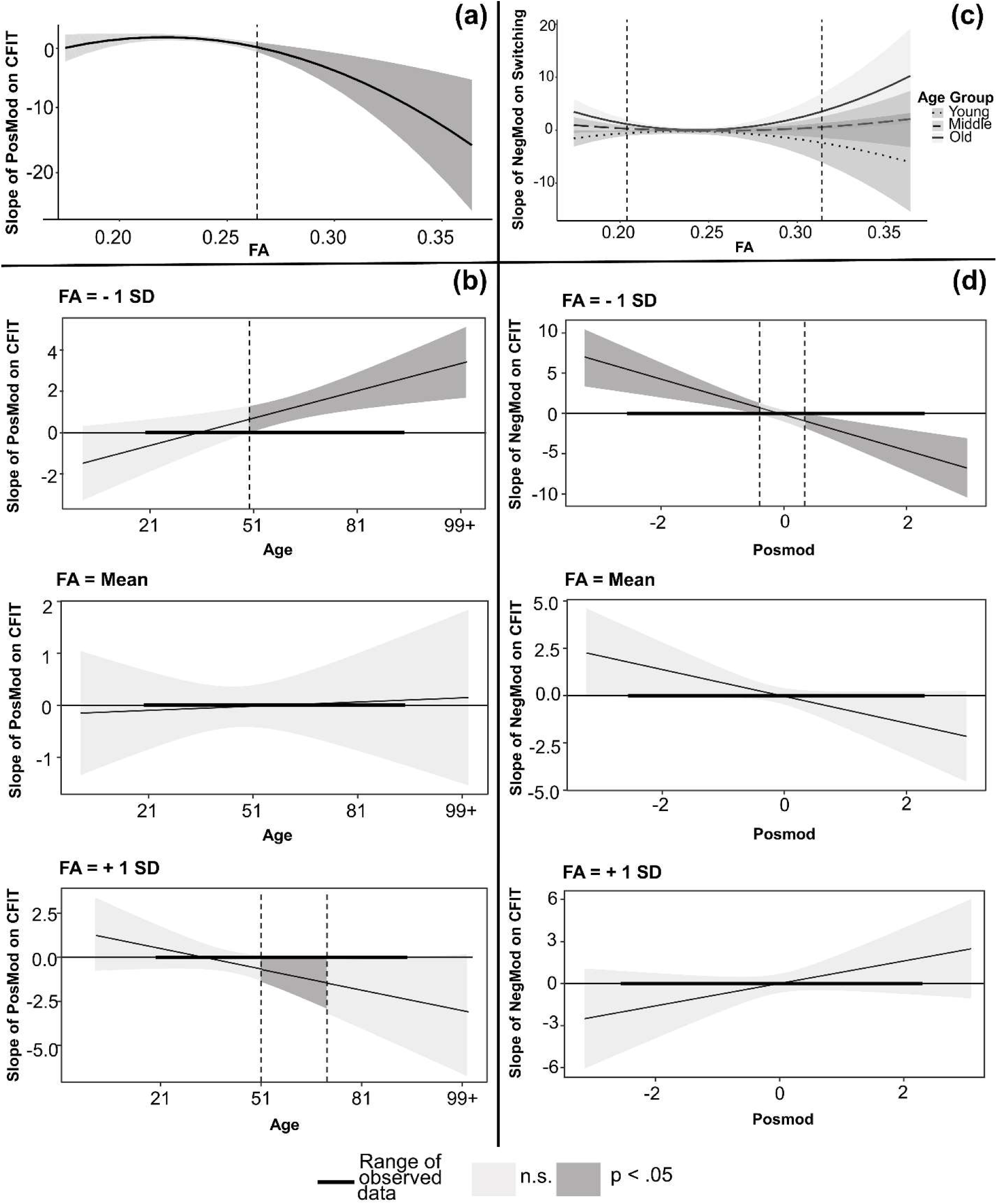
Breakdown of the interactions of the fronto-insular U-fiber tract FA and other independent variables with cognition using Johnson-Neyman plots. **(a)** Interaction of PosMod and quadratic FA on CFIT score **(b)** Interaction of Age, PosMod, and FA on CFIT score, displayed at FA = −1 SD, mean level, and at FA = +1 SD **(c)** Interaction of age, NegMod, and quadratic FA on switching scores **(d)** Interaction of PosMod, NegMod and FA on CFIT scores. FA – fractional anisotropy, CFIT – Culture Fair Intelligence Test; PosMod – positive BOLD modulation to task difficulty, NegMod – negative BOLD modulation to task difficulty.

Within DMN. One *within-network structure-function effect* in the DMN was observed: a significant age × NegMod × Cingulum FA interaction for switching performance (b = 0.66, t(81) = 2.83, *p* = .006, η^2^ = 0.08). Here, age and Cingulum FA modulated the effect of NegMod on switching score. This effect was negative for older adults (> ∼68 years of age) with lower levels of FA, where strengthened NegMod was associated with better switching performance. Nevertheless, a positive effect of NegMod was found for middle-aged and older adults (between ∼55 and 75 years) maintaining relatively high FA. Their strengthened negative BOLD modulation was associated with worse switching performance. See Supplemental Figure 2 and Supplemental Table 3b for detailed statistics.

Between MDN and DMN. We observed a significant 3-way interaction of age × NegMod × U-fiber tract FA^2^ on switching performance (b = −28.68, t(117) = −2.59, *p* = .01, η^2^ = 0.05). This revealed that the relationship between NegMod and switching performance diverged by age, positive for older adults but negative for younger adults. Furthermore, the quadratic interaction indicates that this relationship varied non-linearly across levels of FA within the older adults (+ 1 SD; **Figure 5c** solid line). Specifically, for older adults with higher U-fiber FA, a significant positive association emerged where less negative NegMod was associated with higher switching performance (b = 3.49, t(117) = 2.04, *p* = .04, η^2^ = 0.03). This association was non-significant for older adults at medium levels of FA, but significant for older adults with lower levels of FA (b = 1.12, t(117) = 2.51, *p* = .01, η^2^ = 0.04).

For younger adults (−1SD; **Figure 5c** dotted line), the negative association between NegMod and switching score indicated that strengthened NegMod was associated with better performance. This association was significantly moderated by tract structure, becoming more pronounced at lower levels of FA (b = −0.58, t(117) = −2.26, *p* = .03, η^2^ = 0.03). In other words, for younger adults with lower white matter microstructural integrity in this U-fiber tract, switching performance was more strongly coupled with bold change in DMN regions (NegMod). Meanwhile, there was a non-significant trend for a stronger structure-function association at higher levels of FA for younger adults (b = −2.41, t(117) = −1.29, *p* = .20). The association between Negmod and switching performance was not evident for middle-aged adults (**Figure 5c** dashed line) across any FA level.

#### Coupled network modulation (PosMod × Negmod) and FA

We observed two significant structure-function effects involving coupling of PosMod and NegMod within the U-fiber tract. The first effect was a 3-way interaction of PosMod × NegMod × U-fiber FA on CFIT score (b = 53.22, t(116) = 3.53, *p* < .001, η^2^ = 0.09). The interaction decomposition (via Johnson-Neyman interval analysis; **Figure 5d**) revealed that for participants with relatively low levels of FA, coupling of the PosMod and NegMod was associated with CFIT score. Two types of BOLD modulation coupling were observed: 1) at higher levels of PosMod, the higher the PosMod, the more negative association between NegMod and CFIT score. This aligns with our previous finding (Rieck et al., 2017); 2) At lower levels of PosMod, the association between Negmod and CFIT score shifts, less negative NegMod was associated with higher CFIT scores. This suggests that functional activity in MDN and DMN shifted together when structural integrity is low. Interestingly, these coupling effects were not observed in participants at average or higher levels of U-fiber FA.

We also observed an interaction of age × PosMod × NegMod × U-fiber FA^2^ on inhibition performance (b = 63.68, t(117) = 2.81, *p* = .01, η^2^ = 0.06). This indicates that the joint relationship of PosMod and NegMod with inhibition performance varies as a function of age and FA. (see Supplemental Table 3c). As shown in Figure 6, the interactive patterns across FA levels were relatively comparable between middle-aged and older adults, whereas younger adults exhibited a distinct pattern. For older adults (approximately > 69 years old), the association between NegMod and inhibition score depended on both structural integrity and PosMod. Specifically, at high levels of U-fiber FA (FA > 0.27), paired with high PosMod, less negative NegMod was associated with better inhibition score (b = 2.44, t(117) = 2.07, *p* = .04, η^2^ = 0.03). Within the low-FA subgroup (FA < ∼ 0.18; Figure 6c), a similar pattern was observed where high Posmod is linked with less negative NegMod and better performance (b = 3.37, t(117) = 2.09, *p* = .04, η^2^ = 0.03).

**Figure 6.**
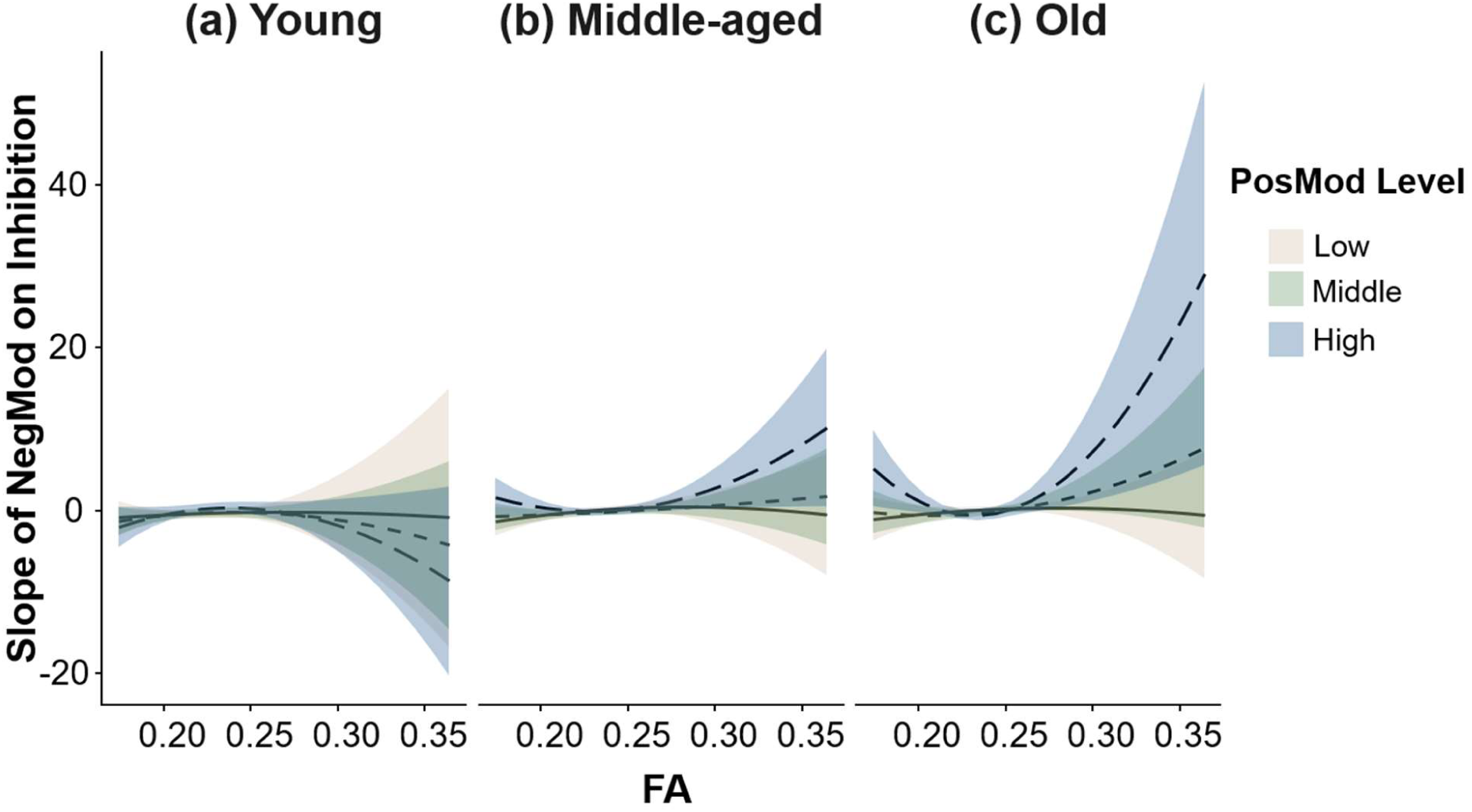
Four-way Interaction of age, PosMod, NegMod, and quadratic FA of the fronto-insular U-fiber tract on Inhibition. Johnson-Neyman plots illustrate the effect of PosMod (at −1, mean and +1 SD), and FA on the slope of NegMod on Inhibition scores at three levels of age (−1, mean, and +1 SD). For older adults, high PosMod (long-dashed line in the right panel c) coupled with low NegMod predicted better Inhibition score when FA is high/low (outside of the range [0.18-0.27]). For middle-aged adults (b), the same coupling effect appeared in higher FA. Quadratic effect of FA in older adults (c) indicates a further weakened NegMod/PosMod coupling effect on inhibiton as FA increases.

Middle-aged adults (around 51 years of age) with high positive BOLD modulation (**Figure 6b**) showed the same coupling effect as older adults at higher levels of FA (b = 1.07, t(117) = 2.09, *p* = .04, η^2^ = 0.03). Still, this coupling effect diminished when FA was relatively average or lower. Younger adults showed a trend for a quadratic effect (inverted U-shape) compared to middle-aged and older adults (U-shape), see **Figure 6a**. In summary, the coupling effect at the heart of the 4-way interaction only appeared in adults middle-aged or older and when frontal-insular U-fiber FA and positive BOLD modulation were relatively high. For older adults with extremely low FA, the coupling effect may have offset the low white matter integrity, as it associated with better inhibition scores. This quadratic effect became weaker with weakened level of PosMod across age groups.

## 4. DISCUSSION

The current study examined the hypothesis that structure-function associations can serve a compensatory role in cognitive aging. To explore these associations, both within- and between networks, we first located four white matter tracts connecting the hub of areas of the multiple demand network (MDN) and, separately, the default mode network (DMN). By incorporating structural and functional brain measures within the same model as predictors of cognitive performance, we were able to observe coupling of positive and negative BOLD modulation as compensatory support of executive function. In accordance with our hypotheses, this coupling effect was observed in middle-aged and older adults, when the white matter in the MDN began to decline. Most importantly, this coupling effect varied dynamically with the microstructural integrity of white matter as indexed by FA.

These significant effects of structure-function association on executive function were most frequently observed for the frontal-insular U-fiber tract, a short-range superficial white matter bundle. Among the four white matter tracts generated by our functionally-guided tractography, the fronto-insular tract was the most vulnerable to age after controlling for the whole brain fractional anisotropy. Additionally, an age-dependent brain structure-function association was observed on another MDN tract (IFOF) and negative BOLD modulation. In contrast to our expectation, older adults with lower MDN FA demonstrated *enhanced* BOLD modulation in the DMN. Below, we discuss potential mechanisms of structure-function compensation across-and within-networks as related to age-related decline, focusing on the U-fiber tract.

### 4.1. Coupling of positive and negative BOLD modulation in compensating for cognitive aging

The motivation for the present study was to explore whether the differential effects of age on structural connectivity and functional activity across the lifespan (i.e., white matter MDN decline occurs before functional changes in that network, but vice versa in the DMN, cross-sectionally) were related to cognitive performance. We hypothesized that weakened negative BOLD modulation might compensate for declined white matter connectivity in the multiple demand network to maintain performance. If the increased BOLD activity is associated with better executive function scores in people with lower white matter connectivity, then we defined it as a compensation effect. The results revealed that weakened negative BOLD modulation compensates for declined white matter in the frontal lobe and reveals a potential reason for “overactivation” in the DMN with increased age. Age-related weakening of negative BOLD modulation or over-activation in DMN regions during task performance has been linked to poorer performance on switching (Basak et al., 2018; Nashiro et al., 2018), *n*-back (Prakash et al., 2012; Qin & Basak, 2020) or executive function tasks (Park et al., 2010; Persson et al., 2007). The weakened negative BOLD activity in response to increasing demand has been considered an example of inefficient deployment of neural resources in adults, associated with reduced cognitive control, and inhibitory deficits (Gazzaley et al., 2005). By contrast, weakened DMN during task performance has also been attributed to greater reliance on prior knowledge in support of goal-directed behavior (Turner & Spreng, 2015). This overactivation in DMN regions across all task difficulty levels would then reflect weakened negative BOLD modulation(Spreng et al., 2014), and could be an indication of a compensatory effect (Duda et al., 2019), or reduced dynamic range of network modulation to increasing task demand (Spreng & Schacter, 2012). The present results partially align with the default-executive coupling hypothesis of aging (DECHA model) (Spreng & Turner, 2019; Turner & Spreng, 2015), which posits that older adults rely on stored existing knowledge or long-term memory representations during task performance. The current findings extend the DECHA model by providing evidence that engagement of default regions can benefit older adults’ cognitive performance.

Previous research demonstrated that coupling of greater BOLD modulation in the MDN and DMN is linked to higher fluid intelligence in the same lifespan sample as that investigated in the present study (Rieck et al., 2017). The present findings extend those prior results by demonstrating that the coupling effect is mainly driven by participants with lower frontal-insular U-fiber tract FA (Figure 5d). For people with lower U-fiber tract FA, better reasoning performance is linked not only to stronger coupling of the two networks but also to the lower BOLD modulation in both networks. Moreover, the combination of MDN engagement and reduced DMN suppression has been linked to older age (Spreng & Turner, 2019; Turner & Spreng, 2015). While functional antagonism between the two networks is associated with cognitive control in healthy adults, age-related decrease in structural connectivity might result in weakened functional activity that lead to either worse behavioral performance or a (compensatory) better performance (Xia et al., 2022). Prior multimodal studies of structural and functional brain aging reported that associations between age and BOLD activation in the DMN were mediated by white matter integrity (Brown et al., 2015; Brown et al., 2018). Further, the same group postulated that age-related decline of white matter might create a noisy environment for functional activation, which decreases the suppression of DMN activity and results in poorer cognitive performance. Similarly, increased functional activity in MDN-related regions may indicate reduced efficiency or failed compensation for structural decline (Brown et al., 2019; Zhu et al., 2015).

In contrast to the previous studies that link functional coupling of the MDN and DMN to poor cognition or decreased behavioral performance, by examining BOLD activity from both networks in the same model (i.e., examining independent and synergistic effects) we were able to show negative coupling of BOLD modulation between the DMN and MDN networks is associated with better executive function in older adults with degraded frontal white matter. One major difference between the current and previous studies is that the current study tested the quadratic effect of white matter integrity in association with these networks, while most studies examined only linear effects. It has consistently been reported that the effects of age on white matter integrity follow a quadratic or cubic function (Lebel et al., 2012; Slater et al., 2019). This nonlinear effect suggests that structural connectivity in MDN might play different roles in modulating the functional activity of the two networks with regard to cognitive performance for older adults. Depending on whether participants possess “high” or “low” FA, the dynamic of this coupling effect changes.

While it is not completely understood how the short-range U-fiber tract connecting inferior frontal regions and insula affects functional activation in MDN and DMN, previous studies have demonstrated the causal influence of the right anterior insula on functional activity in the DMN (Chiong et al., 2013; Sridharan et al., 2008). Similarly, the integrity of the tract connecting the right insula and presupplementary motor area/dorsal anterior cingulate cortex is associated with functional connectivity between right insula and DMN (Jilka et al., 2014). In relation to the present quadratic effect (Figure 9), we propose that age-related decline in frontal U-fiber microstructure is accompanied by alteration in the dynamic coupling of BOLD modulation between the MDN and the DMN. Nevertheless, higher positive and lower negative BOLD modulation were associated with better executive function, and this negative coupling was especially evident in those with relatively high or low FA levels. It is possible that, among middle-aged adults in the early phase of U-fiber degradation, higher FA was associated with an increased capability of utilizing the DMN (increased activation) to compensate for behavioral performance. For older adults in a later phase of U-fiber degradation, those with relatively poor FA also demonstrated the same negative coupling effect, which was not observed among middle-aged adults with poor FA. That being said, the effect of structural integrity on functional activity in aging is not unidimensional, and it may be the case that the involvement of positive and negative BOLD modulation in the early stage of structural decline might act in a compensatory manner, but then later in the aging process show alignment with the notion of “loss of integrity, loss of function” (Andrews-Hanna et al., 2019).

### 4.2. Age-related decrease of white matter microstructure in the multiple demand network

Of the four white matter tracts generated by functionally-guided tractography in the current study, two were MDN-related, one was DMN-related, and one connected both networks (cross-network). Consistent with our hypothesis that white matter connecting regions of the multiple demand system is more vulnerable to increasing age than white matter connecting regions of the default mode system, an age-related reduction in FA of the frontal-insular U-fibers was identified. It is important to note that this effect was above and beyond the general effects of age on global white matter FA. If whole brain FA was not modeled as a covariate of no interest, linear effects of age were observed across all tracts. Contrary to most studies showing a quadratic effect of age on white matter integrity (Lebel et al., 2012), a linear age effect was observed in the current study. One possible explanation is that these tracts are only partial streamlines connecting targeted regions rather than full tracts, and the linear age effect indicates that the streamlines relevant to executive function are susceptible to age as early as their 20s.

However, including whole brain FA in the models is necessary to establish the specificity of the effects to the tracts of interest. The current finding supports previous longitudinal work demonstrating robust aging effects on FA in these short-range white matter tracts (Schilling et al., 2023) and is consistent with our prior finding (Hoagey et al., 2019). In light of the “first in last out” principle (Raz, 2000), it is not surprising to observe age effects on these small tracts, which have a protracted development and reach peak FA later in the lifespan than the longer-range association tracts (Wu et al., 2014). Furthermore, short-range fibers tend to be less myelinated than long-range fibers and hence may be more vulnerable to the deleterious effects of age (Gao et al., 2014).

The structure-function compensation effect within MDN took the form of “over-recruitment” of functional activation in older adults with relatively low white matter FA, where greater BOLD modulation correlated with better performance on inhibition (Figure 6). Reduced white matter integrity in frontal-parietal regions in older adults has been reported to be accompanied by increased BOLD activation or “over-recruitment” during task performance (Burzynska, Garrett, Preuschhof, Nagel, Li, Bäckman, et al., 2013; Daselaar et al., 2015; Zhu et al., 2015), especially in prefrontal cortex. Over-recruitment in older adults, however, has been linked to both better (Burzynska, Garrett, Preuschhof, Nagel, Li, Backman, et al., 2013; Suzuki et al., 2018) or worse (Daselaar et al., 2015; Zhu et al., 2015) executive function. The current results partially account for the previous mixed findings by differentiating MDN- and DMN-related tracts; it appears that the over recruitment - performance association might be dependent on white matter integrity in the MDN. Similarly, the structure-function compensation effect observed in DMN manifested itself as a strengthened BOLD modulation effect (decreasing BOLD signal), which we argue is in compensation for lower white matter integrity (Supplemental Figure 2). Notably, in older adults with degraded cingulum white matter there was a significant inverse association between negative BOLD modulation and switching performance, whereas in adults with higher FA, attenuated BOLD modulation was linked with higher switching scores from middle-age onward. Together, these findings suggest that within-network BOLD modulation might compensate for age-related reductions in white matter integrity.

### 4.3. Cross-network structure-function association

When considering structure-function associations alone, we found that the association between FA and negative BOLD modulation was modulated by age (Figure 4). For younger adults, higher IFOF FA was associated with greater negative BOLD modulation to task difficulty, while older adults demonstrated the opposite association (higher FA correlating inversely with negative modulation). The structure-function association finding in younger adults is consistent with previous studies exploring such associations within the DMN (Brown et al., 2015; Brown et al., 2018; Brown et al., 2019). Additionally, the between-network structure-function association identified here might be explained by the aforementioned “inhibitory” effect of the MDN on the DMN (Chen et al., 2013; Chiong et al., 2013; Sridharan et al., 2008), such that higher structural integrity of the MDN amplified its inhibitory effect on the DMN and, as a consequence, the MDN demonstrated greater negative BOLD modulation with task difficulty. In older adults, however, we observed an opposite structure-function association. Since there was no age-related decline in the integrity of the IFOF tract (after accounting for WBFA), the association identified in older adults might involve modulating factor(s) beyond white matter microstructure, for example, beta-amyloid burden (Foster et al., 2018). Of note, the IFOF tract generated by functionally-guided tractography in the current study connected regions of the MDN (right insula and right posterior parietal), whereas in Brown et al. (2018) the tract was included in an aggregate white matter mask that connected multiple regions of the DMN. Based on its role in deactivation, the IFOF might indirectly interconnect DMN regions, and this indirect connection may contribute to modulation of older adults’ negative BOLD response.

### 4.4 Limitations

The current study was limited, by design, to WM tracts with “direct” connections between two functional clusters, but there might be functionally correlated regions that interact closely without structural connections between them (Hermundstad et al., 2013; Park & Friston, 2013). For example, indirect tracts such as the corpus callosum could be responsible for the interaction of BOLD modulation effects in different brain networks (or hemispheres). Relatedly, the current study focused on the contribution of each of the four white matter tracts connecting BOLD regions within- or across-networks on cognitive aging independently. However, additive or interactive effects in these tracts could impact cognition and will need future studies to explore. The current study is also limited by its cross-sectional design, which only permits identification of age effects on structure-function associations, rather than the effects of aging. The temporal ordering of these hypothesized associations can only be established through the analysis of within-person longitudinal data.

## 5. CONCLUSIONS

The current study utilized functionally-guided tractography derived from task-related functional activations in the multiple demand and default mode networks to examine whether white matter structure – function associations demonstrate compensatory aging effects on cognition, that is, an over recruitment might be linked to better cognitive performance. The results indicated that a single superficial white matter tract (fronto-insular U-fiber tract) in the multiple demand network demonstrated a significant age effect on its microstructure. By examining the quadratic effect of white matter FA, we found evidence that functional effects on executive function and fluid intelligence compensated for low integrity of the U-fiber tract. Consistent with our hypothesis, weakened functional activity in the DMN early in middle age coupled with increased activity in the MDN might represent a compensatory neural response to declining structural connectivity between different regions of the prefrontal cortex.

## DATA AND CODE AVAILABILITY

The data used in the analyses for this paper as well as the R code will be made publicly available on our OSF project page upon publication.

## AUTHOR CONTRIBUTIONS

T-C.L. contributed to data acquisition, project administration, methodology, conceptualization, formal analysis, and manuscript writing, review and editing; D.A.H. contributed to data acquisition, project administration, methodology, and manuscript review and editing; K.M.R. contributed to funding acquisition, supervision, manuscript review and editing; M.D.R. contributed to manuscript review and editing; K.M.K. contributed to funding acquisition, supervision, conceptualization, manuscript review and editing.

## ETHICS

Procedures involving human participants were conducted in accordance with the ethical standards of the 1964 Helsinki Declaration and its later amendments or comparable ethical standards. The study was approved by Institutional Review Boards and all participants provided written informed consent before entering the study.

## ACKNOWLEDGEMENTS

The authors thank Ekarin Pongpipat for assistance with statistical analysis and the research participants for volunteering their time to science.

## Footnotes

1 Because our WBFA variable included voxels from the tracts of interest we tested the models after masking out each tract. Results showed no differences across all the models before or after masking out each tract: Masked WBFA vs WBFA r = .999.

