## Supplementary Materials for "Brain Structure-Function Association as a Compensatory Factor in Cognitive Aging"

**Preliminary tractography**

To examine all possible connections among ROIs and hubs, deterministic tractography was preliminarily applied to all possible ROIs comparisons using less stringent tracking parameters than typical tractography parameters used in previous studies to explore whether any connections were found. After identifying cluster comparisons producing WM connections (i.e., > 0 streamlines produced), deterministic tractography was then applied to each of them with parameters tailored for each tract.

For preliminary tractography, the FA threshold was .05, maximum turning angle of 60°, step size 0 (random selection of 0.5 to 1.5 voxel), a minimum/maximum length of 20/500 mm, no topology-informed pruning (0) was applied, seed orientation started at primary orientation, seed position was subvoxel, smoothing was 0 (propagation direction is independent of previous direction), and all other parameters were set at the default level. Identifying WM streamlines using functionally-defined ROIs is difficult, especially as BOLD clusters are mostly regions of gray matter rather than white matter, and the total number of streamlines of each connection was low. To increase the number of streamlines, these clusters were dilated to include sufficient representation of white matter voxels, using FSL (version 6.0.1). The mean number of white matter voxels within the raw clusters was 1284 voxels, and all clusters were dilated until their white matter voxels exceeded this mean white matter voxel number. It is worth mentioning that the total number of connections (tracts) did not change before and after dilating, only the number of people with streamlines identified, and the total streamlines increased across all connections.

Supp. Table 1. Tracking parameters of tractography

| Tract | Connecting ROIs | Length (mm) | Turning Angle | ROI^a^ | ROA |
| --- | --- | --- | --- | --- | --- |
| IFOF | ROI1-2 | 100-180 | 60 | R Precuneus (AAL) | plane (z = -20) |
| U-fiber | ROI2-4 | 20-50 | 60 |  | plane (z = 20) |
| Cingulum-c | ROI1-8 | 100-150 | 60 | CCsplenium | plane (y= -28) |
| CCsplenium | ROI5-8 | 100-130 | 60 | CC; Cingulum |  |
| Cingulum | ROI8-9 | 130-180 | 50 | Cingulum | ROI5; plane (z = -10) |

a. ROI listed here used additional masks with the clusters to generate the tract; IFG: Inferior frontal gyrus; MFG: Middle frontal gyrus. R: right. ROA: region of avoidance.

Supp. Table 2. Summary of GLMs testing structure-function associations across four tracts. Estimated beta values and *t*-values of main effects and interactions on (a) PosMod and (b) NegMod for each tract.

(a) PosMod

|  |  | β | SE | *t* | *p* |
| --- | --- | --- | --- | --- | --- |
| **Age** | |  |  |  |  |
|  | IFOF | -0.01 | 0.00 | -3.09 | **< 0.01** |
|  | U-fiber | -0.02 | 0.00 | -4.05 | **< 0.01** |
|  | CCsplenium | -0.01 | 0.01 | -2.75 | **0.01** |
|  | Cingulum | -0.01 | 0.00 | -2.97 | **< 0.01** |
| **FA** | |  |  |  |  |
|  | IFOF | -1.59 | 2.10 | -0.76 | 0.45 |
|  | U-fiber | 2.19 | 3.12 | 0.70 | 0.48 |
|  | CCsplenium | -0.83 | 1.71 | -0.49 | 0.63 |
|  | Cingulum | 1.37 | 2.25 | 0.61 | 0.55 |
| **FA^2^** | |  |  |  |  |
|  | IFOF | -83.78 | 41.06 | -2.04 | 0.04 |
|  | U-fiber | -29.25 | 41.13 | -0.71 | 0.48 |
|  | CCsplenium | -3.62 | 23.19 | -0.16 | 0.88 |
|  | Cingulum | 34.76 | 38.51 | 0.90 | 0.37 |
| **WBFA** | |  |  |  |  |
|  | IFOF | 1.28 | 7.36 | 0.17 | 0.86 |
|  | U-fiber | -0.57 | 6.82 | -0.08 | 0.93 |
|  | CCsplenium | -4.85 | 8.72 | -0.56 | 0.58 |
|  | Cingulum | -4.09 | 7.67 | -0.53 | 0.60 |
| **Age * FA** | |  |  |  |  |
|  | IFOF | -0.20 | 0.10 | -1.89 | 0.06 |
|  | U-fiber | -0.16 | 0.13 | -1.27 | 0.21 |
|  | CCsplenium | 0.07 | 0.10 | 0.70 | 0.48 |
|  | Cingulum | -0.01 | 0.11 | -0.11 | 0.91 |
| **Age * FA^2^** | |  |  |  |  |
|  | IFOF | -0.90 | 2.22 | -0.40 | 0.69 |
|  | U-fiber | 3.46 | 3.64 | 0.95 | 0.34 |
|  | CCsplenium | -0.94 | 1.08 | -0.88 | 0.38 |
|  | Cingulum | 1.33 | 2.11 | 0.63 | 0.53 |

a. Bold *p* values indicate significant after correcting for multiple comparison using FDR
b. FA^2^: quadratic effect of FA
(b) NegMod

|  |  | β | SE | *t* | *p* |
| --- | --- | --- | --- | --- | --- |
| **Age** | |  |  |  |  |
|  | IFOF | 0.02 | 0.00 | 4.84 | **< 0.01** |
|  | U-fiber | 0.02 | 0.00 | 4.74 | **< 0.01** |
|  | CCsplenium | 0.01 | 0.00 | 3.44 | **< 0.01** |
|  | Cingulum | 0.01 | 0.00 | 3.59 | **< 0.01** |
| **FA** | |  |  |  |  |
|  | IFOF | 0.31 | 1.74 | 0.18 | 0.86 |
|  | U-fiber | -5.17 | 2.55 | -2.03 | 0.04 |
|  | CCsplenium | -1.99 | 1.32 | -1.51 | 0.13 |
|  | Cingulum | 1.21 | 1.83 | 0.66 | 0.51 |
| **FA^2^** | |  |  |  |  |
|  | IFOF | 69.40 | 34.10 | 2.04 | 0.04 |
|  | U-fiber | 38.37 | 33.67 | 1.14 | 0.26 |
|  | CCsplenium | -3.96 | 17.81 | -0.22 | 0.82 |
|  | Cingulum | -45.20 | 31.22 | -1.45 | 0.15 |
| **WBFA** | |  |  |  |  |
|  | IFOF | 3.59 | 6.11 | 0.59 | 0.56 |
|  | U-fiber | 8.42 | 5.58 | 1.51 | 0.13 |
|  | CCsplenium | 1.53 | 6.70 | 0.23 | 0.82 |
|  | Cingulum | 4.42 | 6.22 | 0.71 | 0.48 |
| **Age * FA** | |  |  |  |  |
|  | IFOF | 0.28 | 0.09 | 3.32 | **< 0.01** |
|  | U-fiber | 0.23 | 0.11 | 2.14 | 0.03 |
|  | CCsplenium | -0.01 | 0.07 | -0.10 | 0.92 |
|  | Cingulum | 0.10 | 0.09 | 1.14 | 0.26 |
| **Age * FA^2^** | |  |  |  |  |
|  | IFOF | -0.48 | 1.85 | -0.26 | 0.80 |
|  | U-fiber | -3.14 | 2.98 | -1.05 | 0.29 |
|  | CCsplenium | -0.24 | 0.83 | -0.29 | 0.77 |
|  | Cingulum | 2.56 | 1.71 | 1.50 | 0.14 |

a. Bold p values indicate significant after correcting for multiple comparison using FDR
b. FA^2^: quadratic effect of FA

Supp. Table 3a. Summary of effects in linear model on accuracy. Effect of Age, PosMod, NegMod, FA, and quadratic FA of four tracts on distance judgment accuracy after controlling for WBFA.

|  | IFOF | | | | U-fiber | | | | CCsplenium | | | | Cingulum | | | |
| --- | --- | --- | --- | --- | --- | --- | --- | --- | --- | --- | --- | --- | --- | --- | --- | --- |
| Model Terms | b | SE | t | p | b | SE | t | p | b | SE | t | p | b | SE | t | p |
| (Intercept) | 0.49 | 0.31 | 1.56 | 0.12 | 0.80 | 0.31 | 2.56 | 0.01 | 0.37 | 0.38 | 0.96 | 0.34 | 0.47 | 0.36 | 1.30 | 0.20 |
| Age | 0.00 | 0.00 | 0.84 | 0.40 | 0.00 | 0.00 | 0.39 | 0.70 | 0.00 | 0.00 | 1.98 | 0.05 | 0.00 | 0.00 | 0.21 | 0.83 |
| PosMod | 0.02 | 0.02 | 1.30 | 0.20 | 0.02 | 0.02 | 1.04 | 0.30 | 0.02 | 0.01 | 1.09 | 0.28 | -0.01 | 0.02 | -0.41 | 0.68 |
| NegMod | -0.07 | 0.02 | -3.94 | **< 0.01** | -0.04 | 0.02 | -2.23 | **0.03** | -0.05 | 0.02 | -2.60 | **0.01** | -0.05 | 0.02 | -2.22 | **0.03** |
| FA | -0.17 | 0.26 | -0.65 | 0.51 | 0.76 | 0.41 | 1.86 | 0.07 | -0.24 | 0.22 | -1.09 | 0.28 | 0.17 | 0.34 | 0.51 | 0.61 |
| FA**^2^** | 15.47 | 5.46 | 2.83 | **0.01** | -3.63 | 9.28 | -0.39 | 0.70 | 5.33 | 3.43 | 1.56 | 0.12 | 0.92 | 6.86 | 0.13 | 0.89 |
| WBFA | 0.90 | 0.78 | 1.15 | 0.25 | 0.16 | 0.79 | 0.21 | 0.84 | 1.25 | 0.96 | 1.30 | 0.20 | 1.01 | 0.90 | 1.12 | 0.27 |
| Age:PosMod | 0.00 | 0.00 | -0.74 | 0.46 | 0.00 | 0.00 | 0.00 | 1.00 | 0.00 | 0.00 | -0.30 | 0.76 | 0.00 | 0.00 | -0.30 | 0.76 |
| Age:NegMod | 0.00 | 0.00 | -1.18 | 0.24 | 0.00 | 0.00 | -0.64 | 0.52 | 0.00 | 0.00 | -0.60 | 0.55 | 0.00 | 0.00 | -0.38 | 0.71 |
| PosMod:NegMod | -0.03 | 0.03 | -1.01 | 0.31 | 0.00 | 0.03 | -0.09 | 0.93 | 0.01 | 0.02 | 0.31 | 0.76 | -0.02 | 0.04 | -0.47 | 0.64 |
| Age:FA | 0.00 | 0.01 | 0.35 | 0.73 | 0.00 | 0.02 | -0.14 | 0.89 | 0.01 | 0.01 | 0.72 | 0.47 | 0.01 | 0.02 | 0.29 | 0.77 |
| PosMod:FA | 0.20 | 0.46 | 0.44 | 0.66 | 0.30 | 0.74 | 0.40 | 0.69 | 0.26 | 0.39 | 0.66 | 0.51 | -0.30 | 0.57 | -0.53 | 0.60 |
| NegMod:FA | 0.21 | 0.46 | 0.46 | 0.65 | 1.20 | 0.73 | 1.65 | 0.10 | 0.75 | 0.46 | 1.61 | 0.11 | -0.74 | 0.52 | -1.42 | 0.16 |
| Age:FA**^2^** | -0.06 | 0.37 | -0.16 | 0.87 | 0.78 | 0.57 | 1.36 | 0.18 | -0.33 | 0.24 | -1.39 | 0.17 | 0.23 | 0.34 | 0.68 | 0.50 |
| PosMod:FA**^2^** | -2.63 | 12.18 | -0.22 | 0.83 | -3.17 | 21.81 | -0.15 | 0.88 | 4.99 | 6.19 | 0.81 | 0.42 | 3.09 | 16.53 | 0.19 | 0.85 |
| NegMod:FA**^2^** | 5.96 | 8.81 | 0.68 | 0.50 | -16.06 | 19.18 | -0.84 | 0.40 | -1.34 | 8.06 | -0.17 | 0.87 | -3.57 | 8.08 | -0.44 | 0.66 |
| Age:PosMod:NegMod | 0.00 | 0.00 | -2.23 | 0.03 | 0.00 | 0.00 | -1.07 | 0.29 | 0.00 | 0.00 | -1.31 | 0.19 | 0.00 | 0.00 | 0.13 | 0.89 |
| Age:PosMod:FA | -0.02 | 0.02 | -0.73 | 0.47 | -0.01 | 0.04 | -0.14 | 0.89 | 0.00 | 0.02 | 0.22 | 0.83 | 0.00 | 0.03 | 0.14 | 0.89 |
| Age:NegMod:FA | 0.01 | 0.02 | 0.30 | 0.76 | -0.05 | 0.04 | -1.16 | 0.25 | 0.03 | 0.03 | 1.02 | 0.31 | -0.02 | 0.03 | -0.76 | 0.45 |
| PosMod:NegMod:FA | -1.87 | 0.76 | -2.47 | 0.02 | 0.01 | 1.12 | 0.01 | 0.99 | 0.88 | 0.71 | 1.24 | 0.22 | -0.03 | 0.94 | -0.04 | 0.97 |
| Age:PosMod:FA**^2^** | 0.00 | 0.73 | 0.01 | 1.00 | -0.50 | 1.18 | -0.43 | 0.67 | -0.08 | 0.42 | -0.18 | 0.86 | -0.76 | 0.81 | -0.94 | 0.35 |
| Age:NegMod:FA**^2^** | -0.18 | 0.58 | -0.31 | 0.76 | -0.85 | 1.25 | -0.68 | 0.50 | -0.27 | 0.55 | -0.50 | 0.62 | -0.93 | 0.54 | -1.73 | 0.09 |
| PosMod:NegMod:FA**^2^** | 41.66 | 17.41 | 2.39 | 0.02 | -34.29 | 43.42 | -0.79 | 0.43 | 8.83 | 16.58 | 0.53 | 0.60 | 19.48 | 35.41 | 0.55 | 0.58 |
| Age:PosMod:NegMod:FA | -0.08 | 0.04 | -2.06 | 0.04 | -0.01 | 0.05 | -0.26 | 0.79 | 0.07 | 0.05 | 1.34 | 0.18 | -0.02 | 0.07 | -0.26 | 0.80 |
| Age:PosMod:NegMod:FA**^2^** | 1.53 | 1.09 | 1.41 | 0.16 | 3.10 | 2.31 | 1.34 | 0.18 | -0.51 | 0.72 | -0.71 | 0.48 | 0.44 | 1.22 | 0.36 | 0.72 |

a. Bold p values indicate significant after correcting for multiple comparison using FDR
b. FA^2^: quadratic effect of FA
c. WBFA: Whole Brain FA

Supp. Table 3b. Summary of effects in linear model on switching score. Effect of Age, PosMod, NegMod, FA, and quadratic FA of four tracts on switching score after controlling for WBFA

|  | IFOF | | | | U-fiber | | | | CCsplenium | | | | Cingulum | | | |
| --- | --- | --- | --- | --- | --- | --- | --- | --- | --- | --- | --- | --- | --- | --- | --- | --- |
| Model Terms | b | SE | t | p | b | SE | t | p | b | SE | t | p | b | SE | t | p |
| (Intercept) | -3.29 | 3.35 | -0.98 | 0.33 | -0.38 | 2.79 | -0.14 | 0.89 | -5.34 | 3.74 | -1.43 | 0.16 | -5.27 | 2.91 | -1.81 | 0.07 |
| Age | -0.02 | 0.01 | -2.60 | **0.01** | -0.01 | 0.01 | -2.53 | **0.01** | -0.01 | 0.01 | -1.71 | 0.09 | -0.02 | 0.01 | -3.06 | **< 0.01** |
| PosMod | 0.21 | 0.19 | 1.09 | 0.28 | 0.21 | 0.15 | 1.43 | 0.15 | 0.16 | 0.14 | 1.08 | 0.28 | 0.07 | 0.15 | 0.48 | 0.63 |
| NegMod | -0.40 | 0.20 | -2.00 | 0.05 | -0.09 | 0.16 | -0.54 | 0.59 | -0.33 | 0.18 | -1.82 | 0.07 | 0.22 | 0.17 | 1.30 | 0.20 |
| FA | 0.79 | 2.79 | 0.28 | 0.78 | 3.21 | 3.65 | 0.88 | 0.38 | -2.03 | 2.15 | -0.94 | 0.35 | -4.08 | 2.72 | -1.50 | 0.14 |
| FA**^2^** | -11.09 | 58.53 | -0.19 | 0.85 | -59.99 | 82.35 | -0.73 | 0.47 | -39.48 | 33.57 | -1.18 | 0.24 | -73.65 | 55.53 | -1.33 | 0.19 |
| WBFA | 8.45 | 8.41 | 1.00 | 0.32 | 1.46 | 6.99 | 0.21 | 0.83 | 13.81 | 9.40 | 1.47 | 0.15 | 13.49 | 7.31 | 1.85 | 0.07 |
| Age:PosMod | 0.01 | 0.01 | 1.07 | 0.29 | 0.01 | 0.01 | 1.29 | 0.20 | 0.00 | 0.01 | 0.14 | 0.89 | 0.00 | 0.01 | -0.46 | 0.65 |
| Age:NegMod | -0.02 | 0.01 | -2.37 | **0.02** | -0.01 | 0.01 | -0.74 | 0.46 | -0.02 | 0.01 | -2.46 | **0.02** | -0.01 | 0.01 | -1.25 | 0.22 |
| PosMod:NegMod | -0.50 | 0.32 | -1.60 | 0.11 | -0.04 | 0.25 | -0.16 | 0.87 | -0.29 | 0.22 | -1.29 | 0.20 | 0.01 | 0.29 | 0.03 | 0.98 |
| Age:FA | -0.01 | 0.14 | -0.06 | 0.95 | 0.28 | 0.19 | 1.50 | 0.14 | 0.09 | 0.13 | 0.70 | 0.49 | 0.02 | 0.15 | 0.14 | 0.89 |
| PosMod:FA | -5.18 | 4.92 | -1.05 | 0.30 | -10.18 | 6.55 | -1.55 | 0.12 | -3.60 | 3.83 | -0.94 | 0.35 | 5.91 | 4.58 | 1.29 | 0.20 |
| NegMod:FA | -4.22 | 4.91 | -0.86 | 0.39 | -5.81 | 6.44 | -0.90 | 0.37 | -4.42 | 4.54 | -0.97 | 0.33 | 5.38 | 4.19 | 1.29 | 0.20 |
| Age:FA**^2^** | -0.83 | 4.01 | -0.21 | 0.84 | -0.13 | 5.08 | -0.02 | 0.98 | 1.63 | 2.36 | 0.69 | 0.49 | -0.80 | 2.72 | -0.29 | 0.77 |
| PosMod:FA**^2^** | -17.68 | 130.52 | -0.14 | 0.89 | -55.77 | 193.45 | -0.29 | 0.77 | -8.10 | 60.62 | -0.13 | 0.89 | 192.74 | 133.79 | 1.44 | 0.15 |
| NegMod:FA**^2^** | 181.78 | 94.41 | 1.93 | 0.06 | 176.29 | 170.10 | 1.04 | 0.30 | 128.54 | 79.01 | 1.63 | 0.11 | -15.77 | 65.37 | -0.24 | 0.81 |
| Age:PosMod:NegMod | -0.03 | 0.02 | -1.80 | 0.07 | -0.03 | 0.02 | -1.72 | 0.09 | -0.03 | 0.01 | -2.17 | 0.03 | 0.00 | 0.01 | 0.24 | 0.81 |
| Age:PosMod:FA | 0.02 | 0.26 | 0.06 | 0.95 | 0.11 | 0.37 | 0.30 | 0.76 | 0.02 | 0.19 | 0.10 | 0.92 | 0.31 | 0.24 | 1.31 | 0.20 |
| Age:NegMod:FA | 0.20 | 0.25 | 0.79 | 0.43 | 0.32 | 0.37 | 0.87 | 0.39 | 0.37 | 0.32 | 1.16 | 0.25 | 0.66 | 0.23 | 2.83 | **0.01** |
| PosMod:NegMod:FA | 4.26 | 8.14 | 0.52 | 0.60 | 3.92 | 9.96 | 0.39 | 0.69 | 1.13 | 6.94 | 0.16 | 0.87 | 0.48 | 7.62 | 0.06 | 0.95 |
| Age:PosMod:FA**^2^** | -7.77 | 7.87 | -0.99 | 0.33 | -16.01 | 10.43 | -1.54 | 0.13 | 3.40 | 4.11 | 0.83 | 0.41 | 9.39 | 6.59 | 1.42 | 0.16 |
| Age:NegMod:FA**^2^** | 1.90 | 6.18 | 0.31 | 0.76 | -28.68 | 11.07 | -2.59 | **0.01** | 1.45 | 5.37 | 0.27 | 0.79 | 3.51 | 4.35 | 0.81 | 0.42 |
| PosMod:NegMod:FA**^2^** | 147.87 | 186.64 | 0.79 | 0.43 | -421.48 | 385.18 | -1.09 | 0.28 | -31.38 | 162.45 | -0.19 | 0.85 | -74.73 | 286.52 | -0.26 | 0.79 |
| Age:PosMod:NegMod:FA | -0.48 | 0.42 | -1.15 | 0.25 | 0.18 | 0.45 | 0.40 | 0.69 | 0.23 | 0.50 | 0.46 | 0.65 | -0.06 | 0.53 | -0.11 | 0.91 |
| Age:PosMod:NegMod:FA**^2^** | 13.62 | 11.63 | 1.17 | 0.24 | 35.61 | 20.50 | 1.74 | 0.08 | 11.24 | 7.01 | 1.60 | 0.11 | -13.10 | 9.90 | -1.32 | 0.19 |

a. Bold p values indicate significant after correcting for multiple comparison using FDR
b. FA^2^: quadratic effect of FA
c. WBFA: Whole Brain FA

Supp. Table 3c. Summary of effects in linear model on inhibition score. Effect of Age, PosMod, NegMod, FA, and quadratic FA of four tracts on inhibition score after controlling for WBFA

|  | IFOF | | | | U-fiber | | | | CCsplenium | | | | Cingulum | | | |
| --- | --- | --- | --- | --- | --- | --- | --- | --- | --- | --- | --- | --- | --- | --- | --- | --- |
| Model Terms | b | SE | t | p | b | SE | t | p | b | SE | t | p | b | SE | t | p |
| (Intercept) | -1.80 | 3.45 | -0.52 | 0.60 | 0.42 | 3.08 | 0.14 | 0.89 | -0.31 | 4.36 | -0.07 | 0.94 | -2.65 | 3.66 | -0.72 | 0.47 |
| Age | -0.02 | 0.01 | -2.76 | **0.01** | -0.02 | 0.01 | -4.19 | **< 0.01** | -0.01 | 0.01 | -1.83 | 0.07 | -0.02 | 0.01 | -2.77 | **0.01** |
| PosMod | 0.10 | 0.20 | 0.50 | 0.62 | 0.14 | 0.16 | 0.84 | 0.40 | 0.18 | 0.17 | 1.06 | 0.29 | 0.05 | 0.19 | 0.24 | 0.81 |
| NegMod | -0.35 | 0.21 | -1.67 | 0.10 | -0.18 | 0.18 | -0.99 | 0.32 | 0.01 | 0.21 | 0.04 | 0.97 | 0.09 | 0.21 | 0.41 | 0.68 |
| FA | 0.51 | 2.88 | 0.18 | 0.86 | 3.73 | 4.03 | 0.92 | 0.36 | -0.12 | 2.51 | -0.05 | 0.96 | -3.68 | 3.42 | -1.08 | 0.28 |
| FA**^2^** | 19.25 | 60.27 | 0.32 | 0.75 | -14.41 | 90.97 | -0.16 | 0.87 | 8.17 | 39.13 | 0.21 | 0.84 | -62.96 | 69.84 | -0.90 | 0.37 |
| WBFA | 4.45 | 8.66 | 0.51 | 0.61 | -0.89 | 7.72 | -0.12 | 0.91 | 0.82 | 10.96 | 0.07 | 0.94 | 6.77 | 9.19 | 0.74 | 0.46 |
| Age:PosMod | 0.00 | 0.01 | -0.25 | 0.80 | 0.00 | 0.01 | 0.55 | 0.59 | 0.01 | 0.01 | 0.90 | 0.37 | 0.00 | 0.01 | -0.16 | 0.87 |
| Age:NegMod | -0.01 | 0.01 | -0.83 | 0.41 | 0.00 | 0.01 | 0.16 | 0.87 | 0.00 | 0.01 | -0.27 | 0.79 | -0.01 | 0.01 | -0.62 | 0.54 |
| PosMod:NegMod | -0.14 | 0.32 | -0.43 | 0.67 | -0.16 | 0.28 | -0.57 | 0.57 | -0.26 | 0.26 | -0.98 | 0.33 | -0.28 | 0.36 | -0.77 | 0.44 |
| Age:FA | -0.21 | 0.15 | -1.44 | 0.15 | -0.05 | 0.21 | -0.27 | 0.79 | -0.03 | 0.15 | -0.22 | 0.83 | -0.12 | 0.19 | -0.66 | 0.51 |
| PosMod:FA | -6.90 | 5.07 | -1.36 | 0.18 | -9.72 | 7.24 | -1.34 | 0.18 | -2.19 | 4.46 | -0.49 | 0.62 | 6.08 | 5.76 | 1.06 | 0.29 |
| NegMod:FA | 2.47 | 5.06 | 0.49 | 0.63 | 10.76 | 7.11 | 1.51 | 0.13 | -1.20 | 5.30 | -0.23 | 0.82 | 6.71 | 5.26 | 1.28 | 0.21 |
| Age:FA**^2^** | 0.78 | 4.13 | 0.19 | 0.85 | 6.67 | 5.62 | 1.19 | 0.24 | -0.09 | 2.75 | -0.03 | 0.98 | 0.33 | 3.42 | 0.10 | 0.92 |
| PosMod:FA**^2^** | -2.55 | 134.41 | -0.02 | 0.98 | 51.00 | 213.70 | 0.24 | 0.81 | -12.16 | 70.65 | -0.17 | 0.86 | 118.23 | 168.27 | 0.70 | 0.48 |
| NegMod:FA**^2^** | 41.34 | 97.23 | 0.43 | 0.67 | 34.60 | 187.91 | 0.18 | 0.85 | -21.96 | 92.08 | -0.24 | 0.81 | -60.52 | 82.22 | -0.74 | 0.46 |
| Age:PosMod:NegMod | 0.00 | 0.02 | -0.17 | 0.86 | -0.03 | 0.02 | -1.90 | 0.06 | -0.01 | 0.01 | -0.70 | 0.49 | -0.01 | 0.02 | -0.38 | 0.70 |
| Age:PosMod:FA | -0.31 | 0.26 | -1.18 | 0.24 | 0.62 | 0.41 | 1.51 | 0.13 | 0.05 | 0.23 | 0.23 | 0.82 | 0.48 | 0.30 | 1.58 | 0.12 |
| Age:NegMod:FA | -0.25 | 0.26 | -0.95 | 0.35 | -0.46 | 0.41 | -1.14 | 0.26 | 0.40 | 0.38 | 1.06 | 0.29 | 0.36 | 0.29 | 1.21 | 0.23 |
| PosMod:NegMod:FA | -5.93 | 8.38 | -0.71 | 0.48 | 4.36 | 11.00 | 0.40 | 0.69 | 3.13 | 8.09 | 0.39 | 0.70 | -2.28 | 9.58 | -0.24 | 0.81 |
| Age:PosMod:FA**^2^** | -1.83 | 8.11 | -0.23 | 0.82 | 1.06 | 11.52 | 0.09 | 0.93 | -4.64 | 4.79 | -0.97 | 0.33 | 2.80 | 8.29 | 0.34 | 0.74 |
| Age:NegMod:FA**^2^** | 4.33 | 6.37 | 0.68 | 0.50 | -15.98 | 12.23 | -1.31 | 0.19 | -4.70 | 6.26 | -0.75 | 0.46 | 2.92 | 5.47 | 0.53 | 0.60 |
| PosMod:NegMod:FA**^2^** | 206.60 | 192.20 | 1.07 | 0.28 | -746.55 | 425.50 | -1.75 | 0.08 | 123.05 | 189.32 | 0.65 | 0.52 | 253.96 | 360.35 | 0.70 | 0.48 |
| Age:PosMod:NegMod:FA | -0.40 | 0.44 | -0.92 | 0.36 | -0.34 | 0.49 | -0.69 | 0.49 | 0.31 | 0.58 | 0.52 | 0.60 | -0.20 | 0.67 | -0.30 | 0.76 |
| Age:PosMod:NegMod:FA**^2^** | -3.90 | 11.98 | -0.33 | 0.75 | 63.68 | 22.65 | 2.81 | **0.01** | 2.22 | 8.17 | 0.27 | 0.79 | 5.38 | 12.46 | 0.43 | 0.67 |

a. Significant p values are marked bold
b. Bold and shading p values indicate significant after correcting for multiple comparison using FDR
c. FA^2^: quadratic effect of FA
d. WBFA: Whole Brain FA

Supp. Table 3d. Summary of effects in linear model on CFIT score. Effect of Age, PosMod, NegMod, FA, and quadratic FA of four tracts on CFIT score after controlling for WBFA

|  | IFOF | | | | U-fiber | | | | CCsplenium | | | | Cingulum | | | |
| --- | --- | --- | --- | --- | --- | --- | --- | --- | --- | --- | --- | --- | --- | --- | --- | --- |
| Model Terms | b | SE | t | p | b | SE | t | p | b | SE | t | p | b | SE | t | p |
| (Intercept) | -0.90 | 4.91 | -0.18 | 0.85 | 2.24 | 4.23 | 0.53 | 0.60 | 3.64 | 5.90 | 0.62 | 0.54 | -2.84 | 4.78 | -0.59 | 0.55 |
| Age | -0.02 | 0.01 | -2.60 | **0.01** | -0.03 | 0.01 | -3.45 | **< 0.01** | -0.03 | 0.01 | -2.56 | **0.01** | -0.03 | 0.01 | -2.97 | **< 0.01** |
| PosMod | 0.32 | 0.28 | 1.14 | 0.26 | 0.62 | 0.22 | 2.81 | **0.01** | 0.37 | 0.22 | 1.67 | 0.10 | 0.21 | 0.25 | 0.86 | 0.39 |
| NegMod | -0.33 | 0.30 | -1.13 | 0.26 | -0.38 | 0.25 | -1.53 | 0.13 | -0.16 | 0.29 | -0.55 | 0.58 | -0.05 | 0.28 | -0.17 | 0.87 |
| FA | -0.38 | 4.11 | -0.09 | 0.93 | 2.64 | 5.53 | 0.48 | 0.63 | 1.00 | 3.40 | 0.29 | 0.77 | -3.48 | 4.45 | -0.78 | 0.44 |
| FA**^2^** | -22.20 | 85.40 | -0.26 | 0.80 | -295.91 | 124.55 | -2.38 | 0.02 | 37.03 | 52.53 | 0.70 | 0.48 | -30.91 | 90.84 | -0.34 | 0.73 |
| WBFA | 19.19 | 12.33 | 1.56 | 0.12 | 11.50 | 10.58 | 1.09 | 0.28 | 7.91 | 14.82 | 0.53 | 0.59 | 24.03 | 11.99 | 2.00 | 0.05 |
| Age:PosMod | 0.01 | 0.01 | 0.78 | 0.44 | 0.01 | 0.01 | 1.15 | 0.25 | 0.02 | 0.01 | 1.56 | 0.12 | 0.01 | 0.01 | 0.79 | 0.43 |
| Age:NegMod | -0.03 | 0.02 | -2.03 | 0.05 | -0.01 | 0.01 | -0.87 | 0.38 | -0.02 | 0.01 | -1.27 | 0.21 | 0.01 | 0.02 | 0.91 | 0.36 |
| PosMod:NegMod | -0.72 | 0.46 | -1.56 | 0.12 | -0.40 | 0.38 | -1.06 | 0.29 | -0.65 | 0.35 | -1.87 | 0.07 | -0.27 | 0.48 | -0.56 | 0.58 |
| Age:FA | -0.22 | 0.21 | -1.06 | 0.29 | -0.75 | 0.28 | -2.67 | **0.01** | 0.09 | 0.20 | 0.46 | 0.65 | 0.08 | 0.24 | 0.33 | 0.74 |
| PosMod:FA | -5.09 | 7.18 | -0.71 | 0.48 | -25.69 | 9.91 | -2.59 | **0.01** | 2.82 | 5.99 | 0.47 | 0.64 | -2.25 | 7.49 | -0.30 | 0.77 |
| NegMod:FA | 9.33 | 7.22 | 1.29 | 0.20 | 4.36 | 9.82 | 0.44 | 0.66 | -3.66 | 7.26 | -0.50 | 0.62 | 7.96 | 6.90 | 1.15 | 0.25 |
| Age:FA**^2^** | -4.20 | 5.86 | -0.72 | 0.48 | 3.80 | 7.70 | 0.49 | 0.62 | -4.57 | 3.70 | -1.24 | 0.22 | -1.37 | 4.46 | -0.31 | 0.76 |
| PosMod:FA**^2^** | -195.76 | 190.39 | -1.03 | 0.31 | -800.22 | 292.83 | -2.73 | **0.01** | -52.74 | 95.00 | -0.56 | 0.58 | 60.30 | 218.89 | 0.28 | 0.78 |
| NegMod:FA**^2^** | -11.28 | 137.91 | -0.08 | 0.93 | 435.42 | 257.37 | 1.69 | 0.09 | -19.56 | 124.27 | -0.16 | 0.88 | -65.15 | 106.97 | -0.61 | 0.54 |
| Age:PosMod:NegMod | -0.01 | 0.02 | -0.61 | 0.55 | -0.04 | 0.02 | -1.81 | 0.07 | -0.01 | 0.02 | -0.65 | 0.51 | 0.02 | 0.02 | 0.73 | 0.47 |
| Age:PosMod:FA | -0.84 | 0.37 | -2.24 | 0.03 | -1.47 | 0.56 | -2.61 | **0.01** | -0.07 | 0.31 | -0.23 | 0.82 | 0.37 | 0.39 | 0.94 | 0.35 |
| Age:NegMod:FA | -0.34 | 0.37 | -0.91 | 0.37 | 0.40 | 0.56 | 0.72 | 0.47 | -0.05 | 0.51 | -0.10 | 0.92 | 0.02 | 0.39 | 0.04 | 0.97 |
| PosMod:NegMod:FA | 9.55 | 11.87 | 0.80 | 0.42 | 53.22 | 15.07 | 3.53 | **< 0.01** | 5.84 | 10.86 | 0.54 | 0.59 | 3.32 | 12.46 | 0.27 | 0.79 |
| Age:PosMod:FA**^2^** | -5.10 | 11.49 | -0.44 | 0.66 | -14.18 | 15.78 | -0.90 | 0.37 | -6.53 | 6.43 | -1.02 | 0.31 | -3.31 | 10.79 | -0.31 | 0.76 |
| Age:NegMod:FA**^2^** | 16.35 | 9.02 | 1.81 | 0.07 | -9.11 | 16.75 | -0.54 | 0.59 | 4.26 | 8.42 | 0.51 | 0.61 | 1.56 | 7.14 | 0.22 | 0.83 |
| PosMod:NegMod:FA**^2^** | 262.03 | 272.22 | 0.96 | 0.34 | -374.90 | 582.76 | -0.64 | 0.52 | 242.10 | 254.13 | 0.95 | 0.34 | -242.38 | 468.91 | -0.52 | 0.61 |
| Age:PosMod:NegMod:FA | 0.16 | 0.62 | 0.27 | 0.79 | 1.28 | 0.68 | 1.89 | 0.06 | -0.60 | 0.78 | -0.76 | 0.45 | -0.25 | 0.87 | -0.29 | 0.77 |
| Age:PosMod:NegMod:FA**^2^** | 5.66 | 16.97 | 0.33 | 0.74 | 72.26 | 31.01 | 2.33 | 0.02 | -8.07 | 10.98 | -0.74 | 0.46 | -10.17 | 16.20 | -0.63 | 0.53 |

a. Bold p values indicate significant after correcting for multiple comparison using FDR
b. FA^2^: quadratic effect of FA
c. WBFA: Whole Brain FA

Supp. Figure1 Johnson-Neyman plots showing quadratic effect of FA of IFOF on Task Accuracy

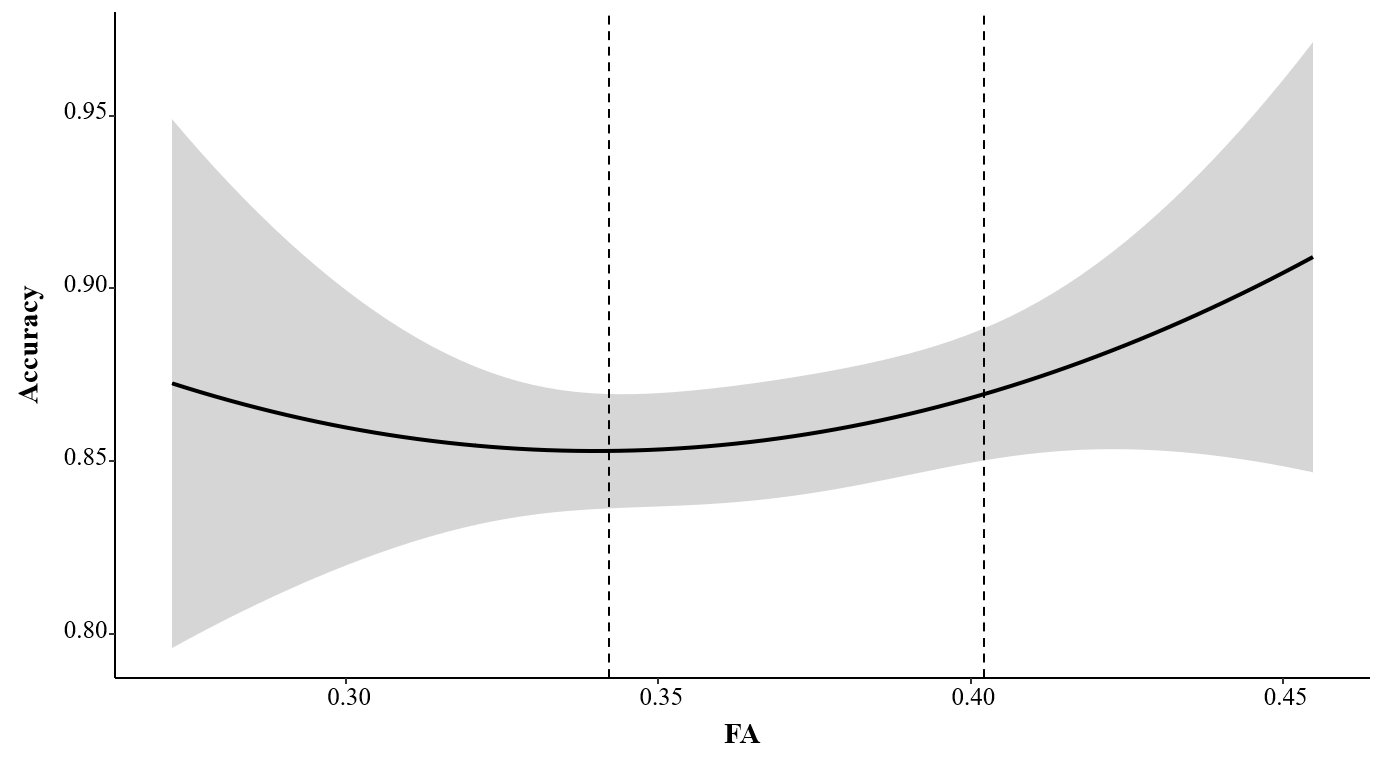

Supp. Figure 2 Johnson-Neyman plots showing Interaction of Age, NegMod, and FA of Cingulum tract on switching score. When FA is low (-1SD), the slope of NegMod on switching is significant during Age older than 67.78; when FA is high (+1SD), the slope of NegMod on switching is significant during Age interval [54.85, 74.89].

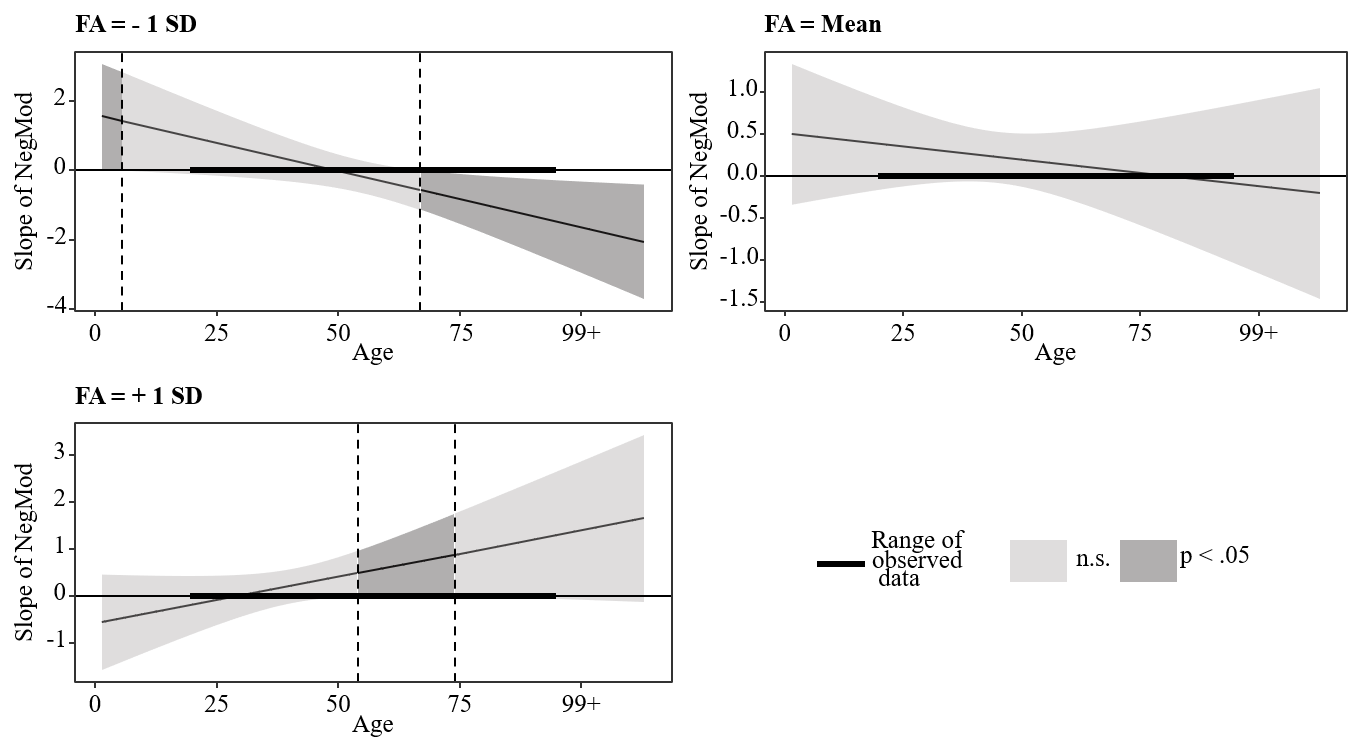
